# Genetic regulation of hepatic retinoid storage is associated with distinct transcriptional networks and liver cellular composition in mice

**DOI:** 10.64898/2026.11.17.767435

**Authors:** Yuta Matsuno, Ayyappa Kumar Sista Kameshwar, Youn-Kyung Kim, Brighton Garrett, Yoshinori Seki, Aashi Srivastava, Marianny Alvarado-Gonzalez, Wellington V. Cardoso, Loredana Quadro, David W. Threadgill, Masako Suzuki

**Author notes:** Address for Correspondence: Masako Suzuki, 498 Olson Blvd, College Station, TX, 77840. Equal contribution (Yuta Matsuno, Ayyappa Kumar Sista Kameshwar).

## Abstract

**Background & Aims:** Although genetic variants influencing circulating retinol (ROH) levels have been reported, the genetic determinants of hepatic vitamin A storage and their biological significance remain poorly understood. We investigated the molecular and cellular mechanisms underlying physiological variation in circulating and hepatic retinoid levels using a genetically diverse inbred mouse population.

**Approach & Results:** Serum ROH, hepatic ROH, and hepatic retinyl ester (RE) concentrations were quantified in 20 inbred mouse strains (n = 4∼5 per strain). Significant strain-dependent variation was observed across all retinoid measures, with heritability estimates of 0.488 for serum ROH, 0.504 for hepatic ROH, and 0.624 for hepatic RE. Integrated liver transcriptomics, untargeted metabolomics, weighted network analyses, and dietary vitamin A intervention studies identified pathways associated with genetically determined retinoid variation. Two hepatic RE-associated transcriptional modules were identified. The metabolic/SMAD4 module was associated with hepatic metabolic pathways and enriched for SMAD4 target genes, whereas the CYP3A module was associated with pathological liver states and disease-associated transcriptional changes. Single-nucleus RNA sequencing revealed strain-dependent differences in liver cell composition. Hepatic RE levels correlated positively with hepatic stellate cell abundance, whereas circulating ROH levels correlated with the proportion of hepatocytes that highly expressed Rbp4.

**Conclusions:** Genetic background significantly influences both circulating retinoid concentrations and hepatic retinoid storage independent of dietary vitamin A intake. Genetic variation shapes retinoid homeostasis through distinct transcriptional programs and differences in adult liver cellular composition, providing new insight into interindividual variation in vitamin A status and susceptibility to metabolic liver disease.

## Introduction

Vitamin A is an essential micronutrient that supports numerous physiological functions and plays a critical role during development^1–4^. Vitamin A status is determined primarily by the dietary intake of animal-derived preformed vitamin A (retinol, ROH, and retinyl esters, RE) and plant-derived provitamin A carotenoids. Upon ingestion, dietary retinoids (vitamin A and its derivatives) are taken up by the enterocytes in the small intestine and packaged as RE into chylomicrons for systemic circulation. Lecithin:retinol acyltransferase (LRAT) is the main enzyme that controls vitamin A esters’ formation in the intestine as well as in other organs. Most dietary retinoids are taken up by the hepatocytes in the liver. Here, RE are hydrolyzed back to ROH, which is subsequently transferred to hepatic stellate cells (HSCs) for storage upon re-esterification^5,6^. The liver is indeed the primary storage site of vitamin A within the body, with the HSC accounting for approximately 50-60% of total body retinoids^7^. As circulating levels of ROH are regulated by homeostasis, liver RE concentration is generally considered the most informative measure of vitamin A status^8^. In times of need, hepatic secretion of ROH bound to retinol-binding protein 4 (RBP4) ensures delivery of vitamin A to the periphery of the body. In the fasting state, RBP4-ROH represents the most abundant form of circulating vitamin A (reviewed by Gottesman et al.^9^). Overall, the liver plays a critical role in maintaining circulating ROH concentrations within a defined range. At the target tissues, cellular uptake of RBP4-ROH can be facilitated by the membrane receptor STRA6^10^. Inside the cell, ROH is enzymatically oxidized first to retinaldehyde and subsequently to retinoic acid (RA), the biologically active form of vitamin A^11^. RA serves as a ligand for retinoic acid receptors (RARs), nuclear transcription factors that regulate the expression of genes involved in development, differentiation, and metabolism. Intracellular RA concentrations -hence its signaling -are tightly regulated not only by its synthesis but also by its degradation mediated by the CYP26 family enzymes^12^.

Although diet is a major determinant of vitamin A status, several lines of evidence, including family studies and genome-wide association studies, indicate that genetic factors also contribute substantially to interindividual variation in circulating retinoid concentrations^13–15^. Moreover, several single nucleotide polymorphisms (SNPs) have been identified as associated with circulating ROH levels (reviewed in ^16,17^). Common genetic variants influencing retinoid metabolism and storage have been identified, including variants in *BCMO1*, which encodes β-carotene 15,15′-dioxygenase responsible for converting provitamin A carotenoids into retinaldehyde, and *PNPLA3*, encoding a lipid hydrolase involved in retinyl ester metabolism^18^ that has also been implicated in hepatic fat accumulation and Metabolic Dysfunction-Associated Steatotic Liver Disease (MASLD) susceptibility^19–23^. However, whether genetic factors that influence circulating retinoid levels also regulate hepatic retinoid storage remains largely unknown.

Elucidating the genetic determinants of systemic vitamin A status in humans is challenging because dietary intake drives much of the variation, and directly assessing hepatic retinoid stores, the body’s primary reservoir of vitamin A, is not feasible in large populations. Although several studies have focused on genetic determinants of circulating retinoid concentrations^14,15,24^, less is known about the factors regulating hepatic retinoid storage. Because the liver is the major reservoir of whole-body retinoids, unraveling the mechanisms that govern hepatic RE accumulation is essential for understanding long-term retinoid homeostasis and its potential role in disease susceptibility. Consequently, genetically defined mouse models provide a valuable system for investigating the genetic regulation of retinoid homeostasis. Obrochta et al. reported strain-dependent differences in serum and hepatic retinoid concentrations among five mouse strains (129S1/SvImJ; C3H/HeJ; BALB/cByJ; AKR/J; C57BL/6J) maintained under different dietary vitamin A conditions^25^. They further demonstrated that the transition from vitamin A-rich to vitamin A-sufficient diets altered retinoid homeostasis in a strain-dependent manner. Despite these observations, the biological significance of physiological variation in hepatic retinoid storage and the mechanisms underlying such variation remain poorly understood.

In this study, we investigated the genetic basis and biological significance of variation in hepatic RE storage by performing multiomics analyses of 100 liver samples from 20 inbred mouse strains, followed by dietary vitamin A supplementation studies on two different common lab inbred mouse strains and single-nucleus RNA sequencing of livers from the most divergent strains with regard to hepatic RE concentrations. We aimed to identify molecular pathways associated with physiological variation in hepatic retinoid storage and determine their potential biological consequences. Given the established roles of retinoid signaling in development and metabolism, we also examined whether variation in hepatic retinoid storage is associated with transcriptional programs related to developmental signaling and metabolic regulation. By defining the genetic and molecular factors that influence hepatic retinoid status, this study provides new insight into how variation in micronutrient status may contribute to differences in disease susceptibility and may ultimately inform precision nutrition and personalized approaches to chronic disease prevention.

## Materials and Methods

### Animal models

The study design is summarized in **Fig. 1A**.

**Fig. 1.**
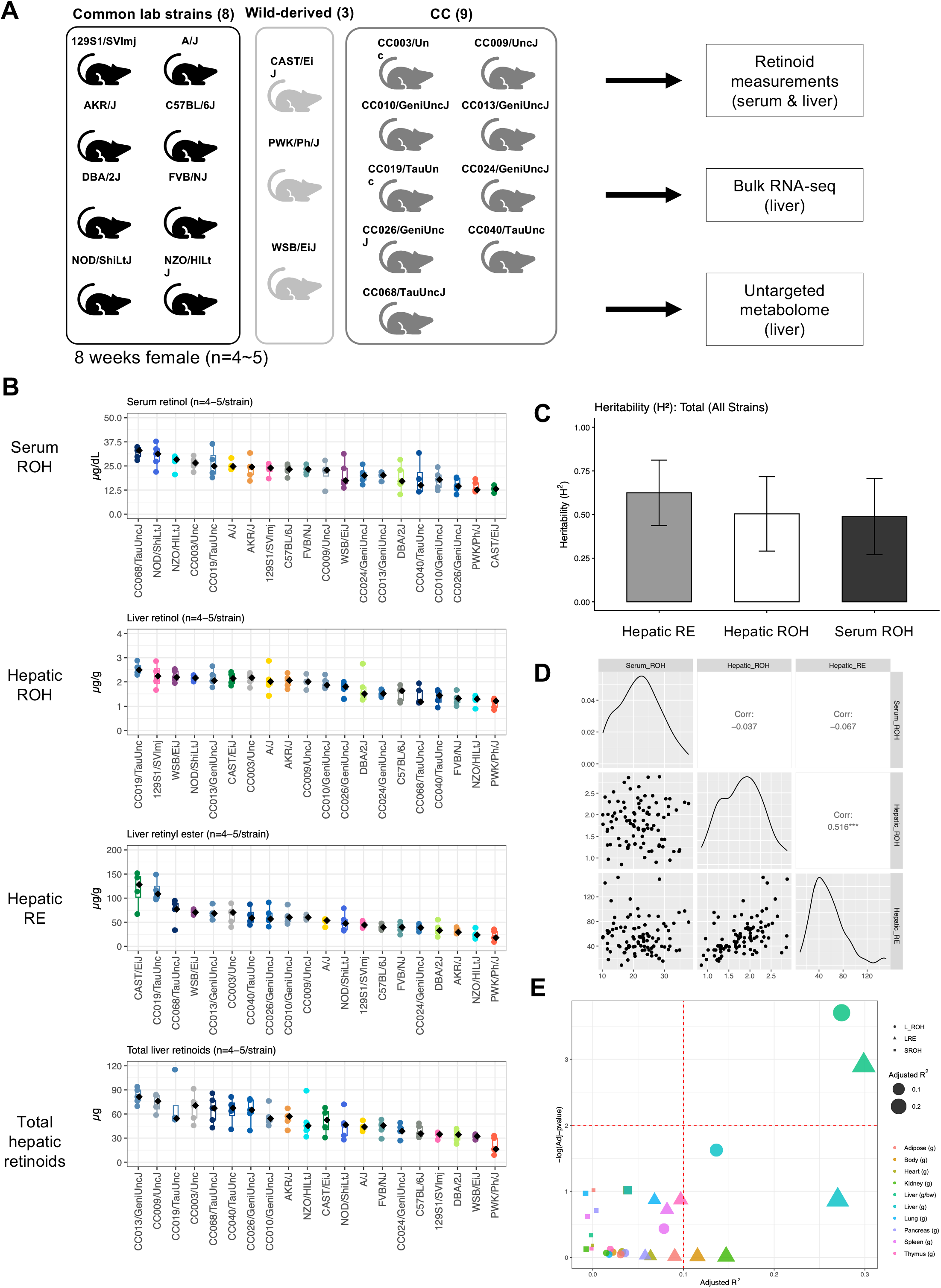
Genetic background determines circulating and hepatic retinoid levels in mice. (A) Experimental design. Twenty genetically diverse inbred mouse strains were analyzed for retinoid phenotypes, liver transcriptomics, and metabolomics. (B) Serum retinol (ROH), hepatic retinol (ROH), hepatic retinyl ester (RE) concentrations, and total hepatic retinoid levels per g of body weight across 20 inbred mouse strains (n = 4-5 per strain). Significant strain-dependent variation was observed for all retinoid measures (one-way ANOVA). (C) Broad-sense heritability (H^2^) estimates for serum ROH, hepatic ROH, and hepatic RE concentrations, demonstrating a substantial genetic contribution to variation in retinoid status. (D) Correlation analysis of serum ROH, hepatic ROH, and hepatic RE concentrations. Hepatic ROH and hepatic RE concentrations were positively correlated, whereas serum ROH was not significantly correlated with either hepatic retinoid measure. (E) Strain-level associations between normalized tissue weights and retinoid measures. Each bubble represents a linear regression model testing the association between a tissue weight trait and serum ROH (square), hepatic ROH (triangle), or hepatic RE (circle) concentrations. Bubble size indicates the adjusted R², and color indicates the tissue. Liver weight normalized to body weight was significantly and negatively associated with both hepatic ROH and hepatic RE concentrations, indicating that strains with smaller livers relative to body weight tended to have higher hepatic retinoid levels. Detailed statistics are provided in **Table S6**.

### Ethics approval

All experiments conducted on mice were approved and carried out in accordance with the ethical guidelines of the Albert Einstein College of Medicine Institutional Animal Care and Use Committee (MS, IACUC# 00001232) and ARRIVE guidelines.

### Inbred mouse strain selection

We selected 20 inbred strains (eight classical laboratory mouse strains, three wild-derived inbreds, and nine Collaborative Cross (CC) recombinant inbreds) for the study. The classical laboratory mouse strains and wild-derived inbreds include eight CC founder strains, 129S1/SVImj (Strain #:002448), A/J (Strain #:000646), C57BL/6J (Strain #:000664), NOD/ShiLtJ (Strain #001976), NZO/HILtJ (Strain #002105), CAST/EiJ (Strain #000928), PWK/Ph/J (Strain #003715), and WSB/EiJ (Strain #038560) in addition to widely used lab strains, AKR/J (Strain #000648), FVB/NJ (Strain #001800), and DBA/2J (Strain #000671). We selected nine CC strains from the 18 phenotypic information available strains (JaxCC1, MPD:519, The Jackson Laboratory Multisystem phenotyping of 18 Collaborative Cross strains (2018), Mouse Phenome Database) based on their body composition and blood chemistry status (blood urea nitrogen, plasma total protein, plasma albumin, plasma calcium, plasma phosphorous, plasma iron, plasma aspartate aminotransferase (AST), plasma alanine aminotransferase (ALT), plasma alkaline phosphatase (ALP), plasma glucose, total cholesterol, high-density lipoprotein (HDL) cholesterol, plasma triglycerides, non-esterified fatty acids (plasma NEFA), total bilirubin, plasma sodium, plasma potassium, plasma chloride, plasma carbon deoxidate, AST/ALT ratio, HDL cholesterol/total cholesterol ratio, and insulin) to sample phenotypically diverse strains (**Fig. S1**). The CC strains used are CC003/Unc, CC009/UncJ, CC010/GeniUncJ, CC013/GeniUncJ, CC019/TauUncJ, CC026/GeniUncJ, CC024/GeniUncJ, CC040/TauUncJ, and CC068/TauUncJ.

### Mice

Seven-week-old female mice were obtained from The Jackson Laboratory (nine classical inbred strains and PWK/PhJ) and the System Genetics Core at the University of North Carolina at Chapel Hill (nine Collaborative Cross [CC] strains). Following a one-week acclimation period, mice were euthanized, and body and tissue weights were recorded. Plasma and liver samples were collected for transcriptomic, metabolomic, and retinoid analyses. During 1-week acclimation, mice were maintained on LabDiet 5LG4 (protein: 22.2%cal, fat: 16%cal, carbohydrate: 61.75%cal, 8.0 IU vitamin A/g), an irradiated diet with ingredients and macronutrient composition comparable to LabDiet 5K52 (protein: 20.1%cal, fat: 16.3%cal, carbohydrate: 61.6%cal, 20.0 IU vitamin A/g), the standard formulation used at The Jackson Laboratory. At the University of North Carolina, CC mice were fed LabDiet® Select Verified 5V0F (protein: 21.0%cal, fat: 15.3%cal, carbohydrate: 63.7%cal, 30.0 IU vitamin A/g). The listed vitamin A concentrations for 5K52 and 5V0F are from the manufacturer’s data sheet before autoclaving. It is known that heat-labile vitamins, such as thiamine (B1), vitamin A, and vitamin E, are reduced after sterilization (autoclaving or irradiation)^26^. In the Kurtz et al. study, the vitamin A concentrations were reduced to about 50-60% after autoclaving.

### Tissue collection

To minimize sample degradation during storage, we dissected the left liver lobe into small pieces (<5 mm), preserved them in RNAlater RNA Stabilization Solution (Invitrogen) for RNA sequencing, and snap-froze them for metabolomic and retinoid analyses. Tissue and plasma samples from CAST/EiJ and WSB/EiJ mice were collected at The Jackson Laboratory and stored as snap-frozen specimens. All tissue and plasma samples were stored at −80°C until analysis. The body and tissue weights summarized by strain are listed in **Table S1**.

### Dietary manipulation

To increase variation in hepatic retinoid status, 8-week-old female C57BL/6J and DBA/2J mice were maintained for 10 weeks on nutritionally matched semi-purified diets containing either 5 or 25 IU vitamin A/g and 10% kcal from fat. Detailed diet compositions are provided in **Table S2**.

### Measuring retinoid concentrations of tissues

Retinol (ROH) and RE were measured by reverse-phase high-performance liquid chromatography (HPLC), as previously reported^27,28^. ROH and RE (retinyl oleate, linoleate, palmitate, and stearate) were identified by comparing the retention times and spectral data of the experimental compounds with those of authentic standards. Retinyl acetate (Sigma, MO, USA) served as an internal standard.

### Broad-sense heritability (H^2^)

H² was estimated for each trait (serum ROH, hepatic RE, and hepatic ROH) with a one-way ANOVA using strain as the grouping factor. The within-strain mean square was used as the environmental variance (Ve), and the genetic variance (Vg) was the between-strain mean square minus the within-strain mean square, divided by the average number of animals per strain. H^2^ is Vg / (Vg + Ve), floored at 0. Error bars show 95% confidence intervals (H^2^ ± 1.96 × SE), truncated at 0 and 1.

### Histopathology study

After tissue dissection, the liver samples were fixed in 4% paraformaldehyde solution at 4°C overnight and processed for paraffin embedding. The paraffin-embedded tissue slides were stained with hematoxylin and eosin (H&E).

### Bulk RNA sequencing (RNA-seq)

#### RNA extraction

After homogenizing tissues with a Bullet Blender Tissue Homogenizer (Speed: 8 and Time: 3 minutes, Next Advance), total RNA was extracted from small pieces of the left liver lobe using the QIAquick RNAeasy plus mini kit (Qiagen, #74134). The quality of the extracted total RNA samples was assessed using the Qubit RNA Integrity and Quality Kit, and the quantity was measured using the Qubit RNA Broad Range Assay Kit. We confirmed that the RNA IQ numbers of the extracted RNA were greater than 8.0 (the average and the standard deviation (SD) were 9.2 and 0.37, respectively). The extracted RNA was stored at -80°C until use.

#### Library preparation, sequencing, and read processing

RNA-seq libraries were generated as non-stranded poly (A) enriched RNA-seq libraries. We sequenced the libraries with 150 paired-end reads, aiming for 6 GB of raw data per sample. The library preparation and sequencing were performed by Novogene, Co (Sacramento, CA). The obtained read sequences were screened using FastQC (version 0.11.9)^29^ to remove low-quality reads, and adapter sequences were eliminated with CutAdapt (version 3.4)^30^. After filtering out low-quality reads and removing adapter sequences, we aligned the sequences to the mouse reference genome GRCm39 with the Gencode basic annotation (version vM36) as annotated transcript information using the STAR aligner (version 2.7.10b) with ENCODE standard options^31^. The number of reads per gene was counted by the STAR aligner’s GeneCounts option. We summarized the sequencing quality statistics in **Table S3**. The average number of sequence reads per sample was 25.6k ± 3.3k paired-end reads, and the average mapping rate was 85.72 ± 2.74% (ranging from 78.29% to 91.08%).

#### Identifying mouse strain-dependently expressed genes and genes associated with retinoid levels

Gene expression counts were normalized for sequencing depth using DESeq2^32^. Genes with low expression (mean count < 2 per sample) were excluded from downstream analyses. Strain-dependent gene expression was evaluated using one-way ANOVA, whereas associations between gene expression and retinoid traits were assessed using linear modeling of the normalized count matrix. Genes with an FDR-adjusted P-value < 0.05 were considered statistically significant.

#### Functional enrichment analyses on identified genes

Functional enrichment analyses were performed on significantly associated genes or module gene sets to identify biological pathways, molecular functions, and regulatory networks represented. The R package Enrichr^33–35^ was used for the analysis. Enriched terms with q-values less than 0.05 were considered significant.

### Quantitative reverse transcription PCR

Total RNA was extracted from samples of the left liver lobe (n=4-5 for each strain), as described above. cDNA was generated using the High-Capacity cDNA Reverse Transcription Kit (Thermo Fisher Scientific, Cat#4374966). Quantitative real-time PCR (qRT-PCR) was performed in duplicate on a LightCycler 480 II (Roche, IN) with SYBR Green Master Mix (Roche Cat# 04707516001). Thermal cycling was programmed for 5 minutes at 95°C for initial denaturation, followed by 45 cycles of 10 seconds at 95°C for denaturation, 20 seconds at 60°C for annealing, and 30 seconds at 72°C for extension and quantification. Primer sets were designed using Primer3, and a single PCR product amplification was confirmed in all assays by melting curve analysis (65 to 95°C). The results were normalized to the housekeeping gene GAPDH. The primers used in this study are listed in **Table S4**.

### Metabolomics analysis

#### Untargeted metabolomics on mouse liver tissues

Untargeted metabolomics analysis was performed on the left liver lobe using ultra-performance liquid chromatography-tandem mass spectrometry (UPLC-MS/MS). The snap-frozen liver samples were homogenized using a ball-mill grinder at 30 Hz for 20 seconds, then metabolites were extracted in 70% methanol. The extracted metabolites were subjected to liquid chromatography (Waters ACQUITY Premier HSS T3 Column 1.8 µm, 2.1 mm * 100 mm) and mass spectrometry (AB Sciex TripleTOF 6600). The untargeted metabolomics analysis was performed by Novogene Co. (Sacramento, CA). To account for technical variation between batches, we used five C57BL/6J liver samples as a reference for the assay.

#### Data processing

Metabolomic data generated in positive and negative ionization modes were merged into a single dataset based on metabolite identifiers and log2-transformed. Batch effects were corrected using a two-step normalization strategy. Within each analytical batch, metabolite abundances were normalized to the median values of the C57BL/6J control samples included in that batch. The normalized datasets were then combined across batches by metabolite identifier. To further minimize residual batch effects, ComBat normalization was performed using the *sva* R package^36^.

#### Strain and retinoid status effects on metabolite abundance in the liver

To assess the effects of hepatic ROH, hepatic RE, and serum ROH on metabolite abundance, a separate linear regression model was fitted for each metabolite. The adjusted R² value was calculated to estimate the proportion of variance explained by each covariate, and the significance of each model was evaluated using the overall F-test. P-values were adjusted for multiple comparisons using the false discovery rate (FDR), and metabolites with FDR-adjusted P values < 0.05 were considered significantly associated with the corresponding covariate.

#### Metabolite set enrichment analysis (MSEA)

Metabolites showing significant associations were subjected to metabolite set enrichment analysis using the overrepresentation analysis (ORA) module of the R package MetaboAnalyst (v4.0)^37–40^, using the corresponding Human Metabolome Database identifiers (HMDB IDs). ORA was performed using the Relational Database of Metabolomic Pathways (RaMP-DB)^41^ as the annotation database. Hypergeometric testing was employed to determine statistical significance, with p-values adjusted for multiple comparisons. An FDR-adjusted p-value less than 0.05 was considered to exhibit significant associations with the pathway.

### Integrated metabolome and transcriptome pathway analysis

We integrated metabolites and genes significantly associated with liver and retinoid levels using MetaboAnalyst R v4.0, with Mus musculus as the reference organism^39^. Gene symbols and HMDB IDs were used to map genes and metabolites to the KEGG database, respectively. Four independent analyses were performed: a) integrated metabolic pathway analysis, which includes pathways containing both metabolites and genes involved in metabolism; b) integrated all-pathway analysis encompassing both metabolic and gene-regulatory pathways; c) metabolite-centric metabolic pathway analysis focusing solely on metabolite enrichment; and d) gene-centric pathway analysis across all pathway types ^42^. All analyses used degree centrality to evaluate network topology and hypergeometric testing to assess pathway enrichment. We applied the “combine queries” approach for metabolite-only (metabolic pathways) and gene-only (all pathways) analyses, which involved direct p-value computation. For the integrated all-pathways and integrated metabolic pathways analyses, we used the “combine p-values” method at the pathway level. We generated network visualizations to depict functional relationships between significant metabolites and genes within enriched pathways, aiding in the mechanistic interpretation of the multi-omics data.

### Processing publicly available liver RNA-seq datasets

Publicly available RNA-seq datasets: Three publicly available liver transcriptomic datasets, GSE72759^43,44^, GSE268273^45^, and GSE193080^46,47^, were included in this study. Raw sequencing data for the mouse dataset (GSE72759) were downloaded from the NCBI Sequence Read Archive (SRA) and processed using the same quality-control, trimming, and alignment procedures described above. For the human datasets, processed and normalized expression data were obtained from GEO and used for subsequent analyses.

### Single-nucleus RNA-seq (snRNA-seq)

Gene expression at single-nucleus resolution was assessed using the 10x Genomics Chromium Single Cell platform on nuclei isolated from frozen liver samples of CAST/EiJ, PWK/PhJ, and WSB/EiJ mice. Libraries were prepared according to the 10x Genomics Chromium Next GEM Single Cell 3’ Gene Expression v4 protocol and sequenced on an Illumina NovaSeq X Plus instrument using a 28-10-10-90 read configuration. Sequencing generated approximately 60 Gb of raw data per library. Raw FASTQ files were processed using Cell Ranger v9.0.1 and aligned to the mouse reference genome 18. Downstream analyses were performed in R using Seurat^48–50^. Nuclei were filtered based on RNA feature and transcript counts (nFeature_RNA = 2,000-7,500; nCount_RNA = 2,000-25,000). Highly variable genes (n = 3,000) were identified, and expression data were scaled prior to integration. Datasets from all strains were integrated using the Seurat data integration workflow, followed by principal component analysis (PCA) on the integrated expression matrix. Uniform Manifold Approximation and Projection (UMAP) was used for dimensionality reduction and visualization. We summarized the sequencing quality and snRNA-seq library statistics in **Table S5**. Cell clusters were annotated using established cell type-specific marker genes ^51,52^ together with results from Seurat’s FindAllMarkers function. Differential gene expression analyses were also performed using the FindMarkers function with the MAST statistical framework. Genes were considered differentially expressed if they met the following criteria: adjusted P < 0.05, |log2 fold change| > 0.58, and expression detected in at least 25% of cells within the cluster.

### Statistical analysis

Statistical analyses were performed using R version 4.4.3 or higher, or GraphPad Prism version 11.1.0 (218). Unless otherwise specified, continuous variables were analyzed using two-tailed Student’s *t-*tests or Wilcoxon Rank-Sum Tests for comparisons between two groups, while Welch’s t-test/ANOVA followed by Tukey’s post hoc test was used for comparisons among more than two groups. P-values less than 0.05, or FDR-adjusted p-values in multiple testing analyses, were considered significant.

## Results

### Genetic background strongly influences hepatic retinoid storage and circulating retinoid levels in mice

Liver and serum RE and ROH were measured by reverse-phase high-performance liquid chromatography (HPLC) as previously described^27^. Significant strain-dependent variation was observed for serum ROH (P =1.34x10^-7^, Welch’s ANOVA), hepatic ROH (P = 6.95 x 10^-8^), hepatic RE (P = 2.55 × 10^-9^), and total hepatic retinoids (p=1.44 x 10^-7^) (**Fig. 1B**). Across strains, serum ROH ranged from 13.1 ± 1.7 μg/dL in CAST/EiJ mice to 32.1 ± 3.1 μg/dL in CC068/TauUncJ mice. Hepatic ROH ranged from 1.13 ± 0.21 μg/g in PWK/PhJ mice to 2.54 ± 0.26 μg/g in CC019/TauUncJ mice, whereas hepatic RE ranged from 22.1 ± 10.7 μg/g in PWK/PhJ mice to 118.7 ± 38.2 μg/g in CAST/EiJ mice. The total hepatic retinoids ranged from 20.79 ± 10.2 μg in PWK/PhJ mice to 82.24 ± 9.85 μg in CC013/GeniUncJ. The highest strain means differed by approximately threefold for serum ROH, fivefold for hepatic RE, and fourfold for total hepatic retinoids.

Interestingly, CAST/EiJ mice exhibited the lowest average circulating ROH concentration and the highest hepatic RE concentration. However, this inverse relationship was not consistently observed across strains, suggesting that circulating and stored retinoid pools are regulated at least partially independently. PWK/PhJ mice showed the lowest averages in hepatic ROH, RE, and total retinoids (**Fig. 1B**). Detailed post hoc comparisons are provided in **Table S6**.

Broad-sense heritability (H²) was estimated at 0.488 for serum ROH, 0.504 for hepatic ROH, and 0.624 for hepatic RE, indicating a substantial genetic contribution to variation in vitamin A status (**Fig. 1C**). Broad-sense heritability estimates did not differ significantly among classical laboratory, wild-derived, and CC strain groups (**Fig. S2A**). Although comparable estimates for hepatic RE are unavailable in humans, the heritability values observed here exceeded those reported for circulating ROH in human studies^13^, likely reflecting the reduction in environmental variation achieved by maintaining all animals under comparable standard diet conditions.

Consistent with the distinct regulation of circulating and hepatic retinoid pools, hepatic RE and hepatic ROH concentrations were positively correlated (r = 0.516), whereas serum ROH was not significantly correlated with either hepatic retinoid measure (**Fig. 1D**, **Fig. S2B**). Together, these findings demonstrate that both circulating retinoid levels and hepatic retinoid storage are strongly influenced by genetic background and are consistent with complex genetic regulation of retinoid homeostasis.

We next examined the relationship between retinoid status and major tissue weights (heart, kidney, liver, lung, pancreas, spleen, and thymus) and body weight. Liver weight normalized to body weight was negatively associated with both hepatic RE levels (β = −15.23 wt%, adjusted R² = 0.299, adjusted P = 0.00124) and hepatic ROH levels (β = −0.246 wt%, adjusted R² = 0.274, adjusted P = 0.000198), indicating that strains with smaller livers relative to body size tended to exhibit higher hepatic retinoid concentrations (**Fig. 1E**). Histopathological assessments using H&E staining among CC strains showed no significant variation associated with hepatic RE or hepatic ROH levels (**Fig. S3**).

### Genetic background influences hepatic gene expression and metabolite profiles, uncovering genes and metabolites associated with retinoid levels

To investigate molecular mechanisms underlying physiological variation in vitamin A status, we performed bulk RNA-seq and untargeted metabolomics on the same liver samples used for retinoid measurements (n = 4-5 per strain; 20 inbred strains). Principal component analysis of the transcriptomic data revealed highly strain-specific gene expression profiles despite comparable standard dietary conditions (**Fig. 2A**). Strain was the dominant determinant of liver transcriptional variation, explaining 52.4% of the variance in PC1 (P < 0.001). Although hepatic ROH and hepatic RE were also significantly associated with PC1, they explained much less than that of strain (0.4% and 3.8%, respectively) (**Fig. 2B**). No other evaluated variables were significantly associated with PC1 variation. The percentage of uniquely mapped reads was not significantly associated with gene expression variation, indicating minimal technical influence on transcriptomic profiles. Consistent with this conclusion, qRT-PCR validation demonstrated strong concordance with RNA-seq results (**Fig. S4A**).

**Fig. 2.**
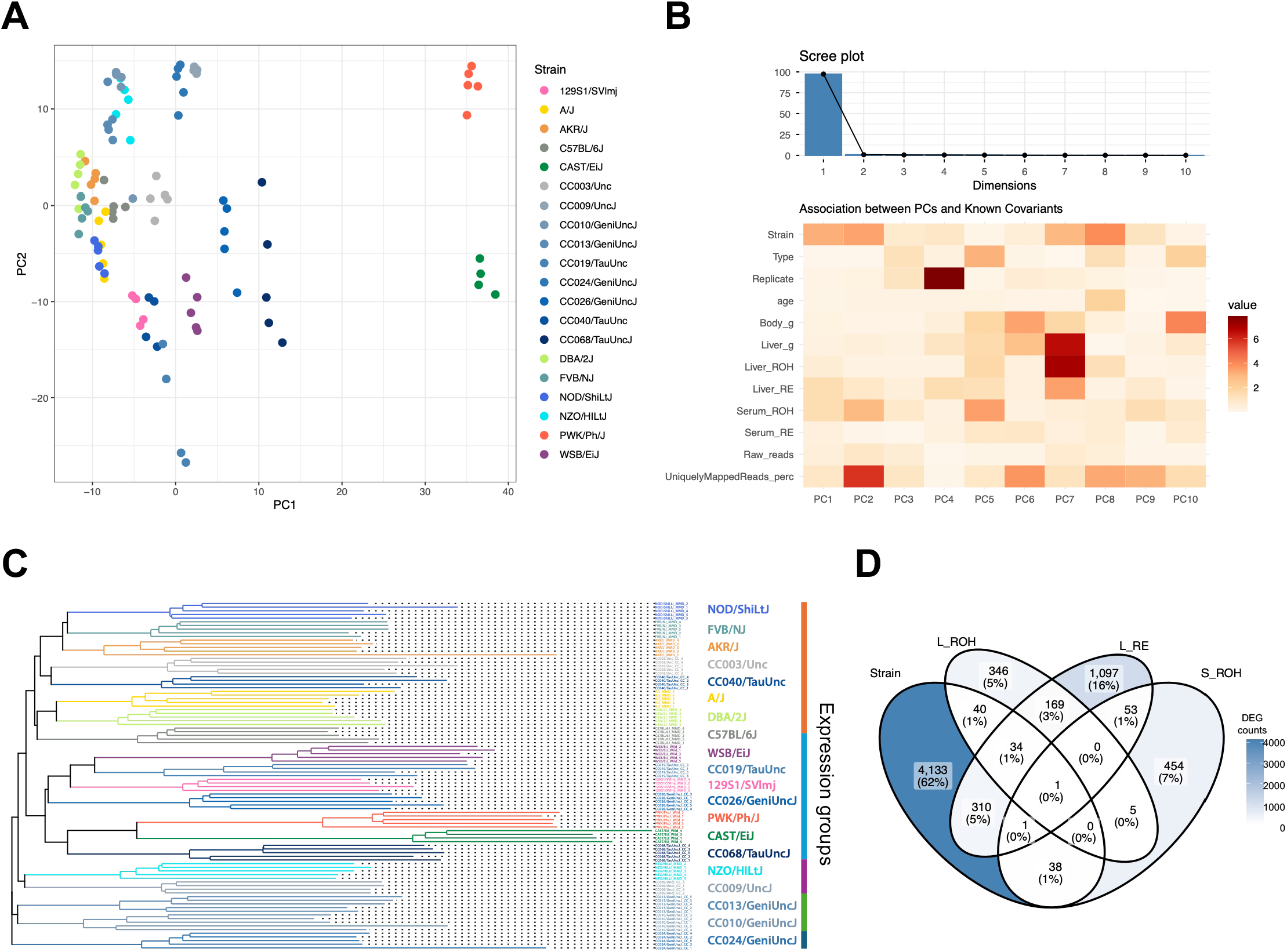
Genetic background influences hepatic gene expression profiles. (A) Principal component analysis of liver transcriptomic data revealed distinct strain-specific gene expression profiles across 20 inbred mouse strains. (B) Association analysis of factors contributing to transcriptional variation in liver RNA-seq data. Strain was the dominant contributor to PC1 variation, explaining 52.4% of the variance (P < 0.001). Although hepatic ROH and hepatic RE were also significantly associated with PC1, they explained substantially less variance (0.4% and 3.8%, respectively). No other evaluated variables were significantly associated with PC1 variation. (C) Hierarchical clustering of liver transcriptomes across 20 inbred mouse strains. Five major transcriptional clusters were identified, although clustering patterns did not fully correspond to strain phylogenetic relationships. (D) Overlap between genes exhibiting significant strain-dependent expression and genes associated with hepatic retinyl ester (RE), hepatic retinol (ROH), or serum ROH concentrations. Hepatic RE-associated genes showed the greatest overlap with strain-dependent genes (345 genes), compared with hepatic ROH-associated genes (74 genes) and serum ROH-associated genes (39 genes).

Hierarchical clustering further demonstrated pronounced strain-dependent transcriptional signatures and identified five major gene expression clusters (**Fig. 2C**). Notably, clustering patterns only partially reflected known phylogenetic relationships among strains, suggesting that transcriptional variation captured biological differences beyond genetic relatedness alone. Differential expression analyses identified 4,565 genes exhibiting significant strain-dependent expression, as well as 1,669, 596, and 554 genes significantly associated with hepatic RE, hepatic ROH, and serum ROH levels, respectively (**Table S7**). Among these, 345 hepatic RE-associated genes, 74 hepatic ROH-associated genes, and 39 serum ROH-associated genes also exhibited significant strain-dependent expression (**Fig. 2D**, **Fig. S4B**).

Functional enrichment analysis revealed distinct biological processes associated with each retinoid measure. Genes positively associated with hepatic ROH were enriched for fatty acid metabolism and insulin response pathways (**Fig. S5**), whereas serum ROH-associated genes were enriched for protein stability and mitochondrial functions (**Fig. S6**). In contrast, genes positively associated with hepatic RE were strongly enriched for metabolic pathways, including glycerolipid, glycerophospholipid, phospholipid, and glucose metabolism (**Fig. S7**). These findings suggest that physiological variation in hepatic RE storage is closely associated with strain-specific metabolic programs.

To determine whether physiological variation in hepatic retinoid status is also reflected at the metabolomic level, we performed untargeted metabolomics on the same liver samples. A total of 2,107 metabolites were detected. Like the transcriptomic analyses, principal component analysis revealed strong strain-dependent metabolic signatures (**Fig. 3A**). However, substantially fewer metabolites than genes were associated with retinoid measures (**Fig. S8**; **Table S7; S8**).

**Fig. 3.**
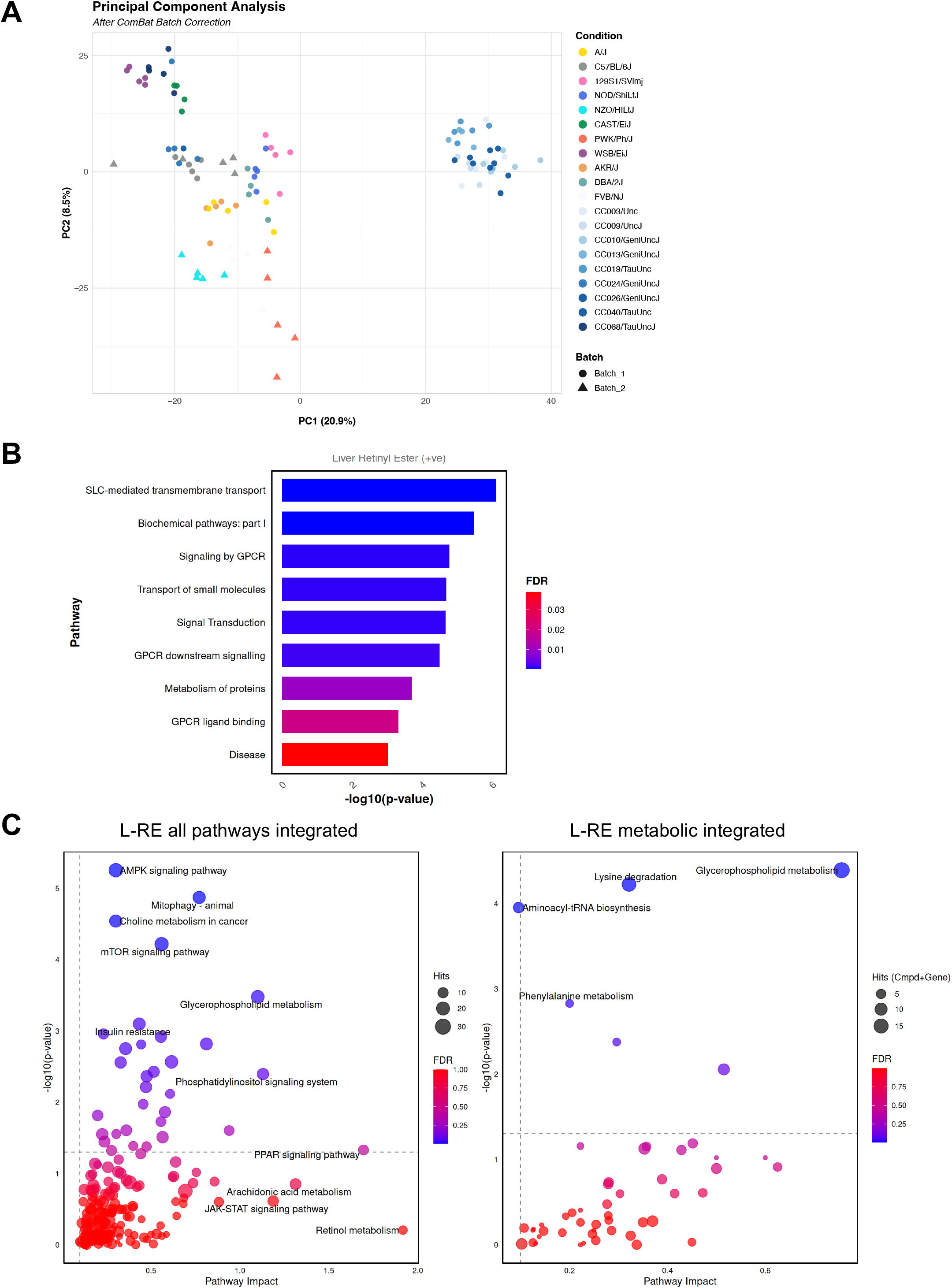
Genetic background influences hepatic metabolomic profiles and identifies metabolic pathways associated with hepatic retinoid storage. (A) Principal component analysis of liver metabolomic profiles. Similar to the transcriptomic data, samples clustered primarily according to strain, indicating strong strain-dependent metabolic signatures. (B) Metabolite set enrichment analysis of metabolites positively associated with hepatic retinyl ester (RE) concentrations. Hepatic RE-associated metabolites were enriched in G protein-coupled receptor (GPCR) signaling pathways. (C) Integrated pathway analysis of genes and metabolites associated with hepatic RE levels. Left, enrichment analysis using all KEGG pathways, including metabolic and gene-only signaling pathways. Right, enrichment analysis restricted to KEGG metabolic pathways containing both metabolites and metabolic genes. Pathway significance was determined using MetaboAnalyst joint pathway analysis.

Metabolite set enrichment analysis showed that metabolites positively associated with hepatic RE were enriched in GPCR signaling-related pathways, whereas hepatic ROH-associated metabolites were enriched in transporter and amino acid metabolism pathways (**Fig. 3B**, **Fig. S9A**). No enriched pathways were identified for serum ROH-associated metabolites.

To identify biological pathways associated with hepatic retinoid storage, we performed integrated metabolomic-transcriptomic pathway analysis using MetaboAnalyst. This analysis was conducted using two reference pathway sets: (i) all KEGG pathways, which include both metabolic and gene-only signaling pathways, and (ii) metabolic pathways only, which are limited to pathways containing both metabolites and metabolic genes. When all KEGG pathways were considered, more than 25 pathways were significantly associated with hepatic RE levels, including glycerophospholipid metabolism, insulin resistance, mTOR signaling, PPAR signaling, and choline metabolism in cancer (**Fig. 3C, left; Fig. S10**). In contrast, restricting the analysis to metabolic pathways identified a smaller set of pathways jointly supported by both metabolite and gene alterations, with glycerophospholipid metabolism emerging as the most significantly enriched pathway, followed by lysine degradation, aminoacyl-tRNA biosynthesis, and phenylalanine metabolism (**Fig. 3C, right**). In contrast to hepatic RE, only two pathways were significantly associated with hepatic ROH (**Fig. S9B**), suggesting that pathway-level metabolic and regulatory changes are more strongly linked to hepatic RE levels.

### WGCNA and WMCNA identified gene and metabolite modules associated with genetically determined hepatic retinoid status, and module correlation analysis revealed associations between these molecular networks

To identify molecular networks associated with physiological variation in hepatic retinoid status, we performed Weighted Gene Correlation Network Analysis (WGCNA) using retinoid levels as traits of interest. WGCNA identified 23 gene co-expression modules, of which two modules (brown and blue) were positively associated with hepatic RE levels and one module (midnight blue) was negatively associated with hepatic RE levels (**Fig. 4A**). The brown module contained 1,267 genes, whereas the blue module contained 761 genes (**Table S9**). Consistent with the association analyses shown in **Fig. 1E**, liver weight normalized to body weight was inversely associated with the RE-associated gene modules, suggesting a relationship among hepatic RE levels, retinoid-associated transcriptional programs, and relative liver mass.

**Fig. 4.**
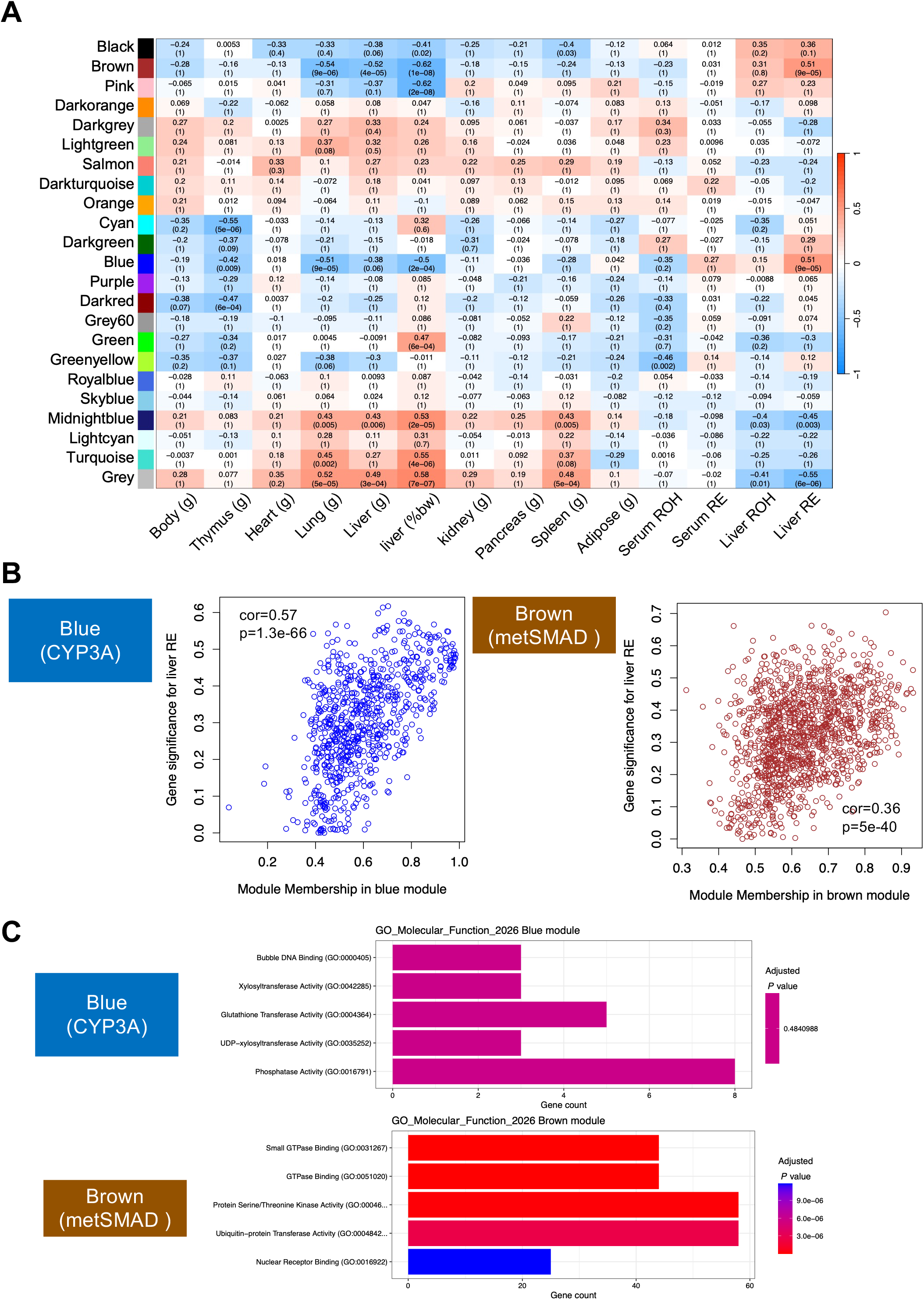
WGCNA identifies hepatic retinoid-associated gene co-expression modules. (A) Module-trait relationship heatmap from weighted gene correlation network analysis (WGCNA). Two modules (brown and blue) were positively associated with hepatic retinyl ester (RE) levels, whereas the midnight blue module was negatively associated with hepatic RE levels. (B) Relationship between module membership (MM) and gene significance (GS) for hepatic RE levels in the brown and blue modules. In both modules, genes with higher intramodular connectivity exhibited stronger associations with hepatic RE levels, indicating that highly connected hub genes were strongly associated with hepatic retinoid storage. (C) Gene Ontology (GO) enrichment analysis of genes within hepatic RE-associated gene modules. Top, no significantly enriched GO terms were identified for the blue module. Bottom, the brown module was enriched for GO terms related to GTPase binding, protein serine/threonine kinase activity, and ubiquitin-protein transferase activity.

We next focused on the two positively associated modules. In both modules, module membership was positively associated with gene significance for hepatic RE levels, indicating that highly connected hub genes were also those most strongly associated with hepatic retinoid storage (**Fig. 4B**). Functional enrichment analysis revealed no significant Gene Ontology (GO) terms for the blue module (**Fig. 4C, Fig. S11**). In contrast, the brown module was enriched for GO terms related to GTPase binding, protein serine/threonine kinase activity, and ubiquitin-protein transferase activity (**Fig. 4C, Fig. S12**). In addition, transcription factor enrichment analysis revealed significant enrichment of SMAD4-binding motifs among brown module genes (**Fig. S13A**). Hepatic *Smad4* expression was positively associated with hepatic retinyl ester (RE) levels (adjusted R² = 0.1385, P = 0.0012; **Fig. S13B**), and a large proportion of brown module genes exhibited significant associations with *Smad4* expression (**Fig. S13C**). Together, these findings identify SMAD4 as a candidate upstream regulator of the brown module and implicate SMAD4-associated transcriptional programs in physiological variation in hepatic retinoid storage. Among the highly connected blue-module hub genes, Cyp3a25 showed strong associations with retinoic acid receptor genes (RARs and RXRs) (**Fig. S14**). Accordingly, we hereafter refer to the blue module as the CYP3A-associated (CYP3A) module. The metSMAD module was enriched for SMAD4-regulated genes and metabolic functions, whereas the CYP3A module was characterized by CYP-family genes, including *Cyp3a25*.

Weighted Metabolite Correlation Network Analysis (WMCNA) identified eight metabolite modules, of which the brown module was significantly and positively associated with both hepatic RE and ROH levels (**Fig. 5AB**). The brown module contained 277 metabolites, 141 of which had HMDB identifiers and were used for pathway enrichment analysis. These compounds were significantly enriched in several chemical classes and subclasses, including amino acids, peptides, and analogs; fatty acids and conjugates; carbohydrates and carbohydrate conjugates; and bile acids, alcohols, and derivatives (**Fig. S15AB**). Pathway enrichment analysis showed that metabolites in this module were enriched for GPCR signaling pathways (**Fig. 5C**), further supporting the regression analyses of metabolites that implicated GPCR signaling as a pathway associated with hepatic retinoid status (**Fig. 3B**). Enriched Metabolite Sets analysis using dbGaP identified significant enrichment for metabolites associated with the single-nucleotide polymorphism (SNP) rs1532085 (**Fig. S15C**). rs1532085 is located upstream of the human hepatic lipase gene (*LIPC*), and its genotype has been reported to be associated with serum lipid parameters^53^. Furthermore, GTEx data indicate that rs1532085 is associated with the expression levels of *LIPC* and Aldehyde dehydrogenase 1 family, member A2 (*ALDH1A2*), suggesting a potential regulatory role for this variant (**Fig. S15D**). While no associations between *LIPC* and retinoid status have been reported, *ALDH1A2* codes an enzyme that catalyzes the synthesis of all-trans retinoic acid (RA) from retinaldehyde^54^. Thus, this observation raises the possibility that retinoid signaling contributes to the metabolic signatures associated with the brown metabolite module.

**Fig. 5.**
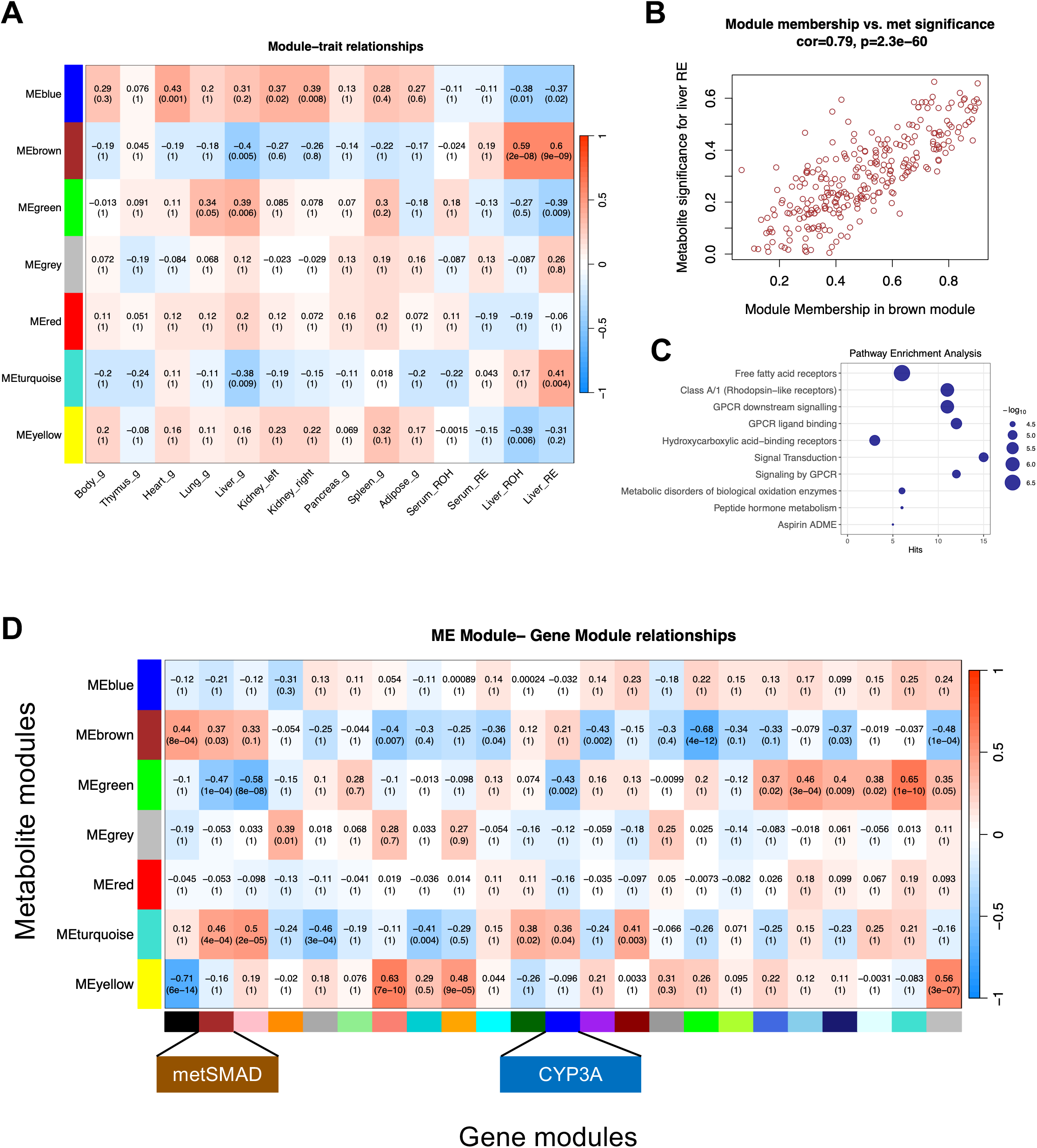
WMCNA identifies hepatic retinoid-associated metabolite modules. (A) Module-trait relationship heatmap from weighted metabolite correlation network analysis (WMCNA). Among the eight metabolite modules identified, the brown module was positively associated with both hepatic retinyl ester (RE) and hepatic retinol (ROH) concentrations. (B) Relationship between metabolite module membership (MM) and metabolite significance (MS) for hepatic RE levels in the brown module. Metabolites with higher intramodular connectivity exhibited stronger associations with hepatic RE levels, indicating that highly connected metabolites were associated with hepatic retinoid storage. (C) Pathway enrichment analysis of metabolites within the brown metabolite module. Metabolites in this module were enriched for G protein-coupled receptor (GPCR) signaling pathways. (D) Association analysis between WGCNA gene modules and WMCNA metabolite modules. The metSMAD gene module was positively associated with the brown metabolite module (r = 0.37, FDR-adjusted P = 0.03), indicating coordinated variation between hepatic gene expression and metabolite profiles associated with retinoid storage. In contrast, the CYP3A module was not significantly associated with the brown metabolite module.

Module association analysis between WGCNA gene modules and WMCNA metabolite modules revealed a significant positive association between the metSMAD module and the brown metabolite module (r = 0.37, FDR-adjusted P = 0.03), indicating coordinated variation between hepatic gene expression and metabolic profiles associated with retinoid storage (**Fig. 5D**). In contrast, the CYP3A module was not significantly associated with the brown metabolite module (FDR-adjusted P = 1.00), suggesting that its transcriptional variation is largely independent of the identified metabolic network (**Fig. 5D**). Together, these findings indicate that the metSMAD and CYP3A modules represent distinct molecular programs associated with hepatic retinoid status, with the metSMAD module more closely linked to metabolic variation in the liver.

### Gene expression modules associated with genetically determined hepatic retinoid status are largely independent of those associated with dietary retinoid perturbations

Dietary vitamin A is the primary source of retinoids and plays a central role in regulating tissue and circulating retinoid levels. Although source site and dissection site were not associated with retinoid levels, transcriptomic profiles, or metabolomic profiles, and all mice were maintained on LabDiet 5LG4 for one week before tissue collection (except CAST/EiJ and WSB/EiJ), the strains were obtained from two different vendors and had been maintained on chow diets containing different amounts of vitamin A prior to study. Thus, residual effects of pre-study dietary exposure cannot be completely excluded. To determine whether dietary vitamin A influences gene expression within the metSMAD and CYP3A modules, we performed a controlled dietary intervention study using female C57BL/6J and DBA/2J mice. These widely used laboratory strains did not differ significantly in serum ROH, hepatic ROH, hepatic RE, or total hepatic retinoid concentrations under standard chow conditions (Tukey’s HSD-adjusted P > 0.9999; **Table S6**) and were therefore selected to evaluate the effects of dietary vitamin A independent of baseline differences in retinoid status. Mice were fed diets containing either 5 or 25 IU vitamin A/g for 10 weeks (n = 5 per strain per diet group). The 5 IU vitamin A/g diet approximates the vitamin A level recommended in the AIN-93 formulation, whereas the 25 IU vitamin A/g diet provides a substantially higher amount comparable to that present in standard rodent chow. Hepatic retinoid concentrations were measured, and liver transcriptomes were profiled by bulk RNA-seq (**Fig. 6A**).

**Fig. 6.**
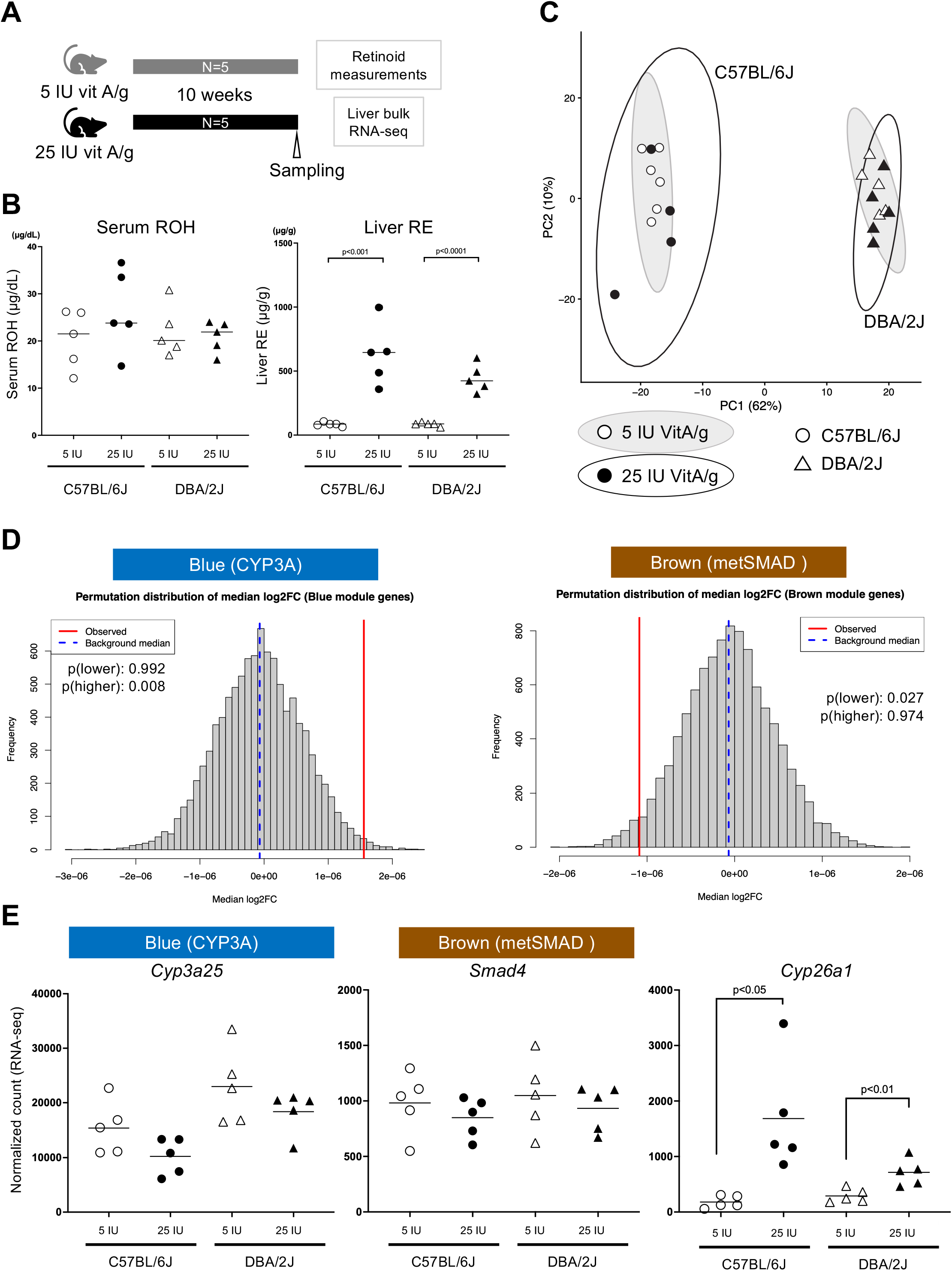
Hepatic retinoid-associated gene modules are largely independent of dietary vitamin A supplementation. (A) Experimental design for the dietary vitamin A supplementation study. Female C57BL/6J and DBA/2J mice were fed diets containing either 5 or 25 IU vitamin A/g for 10 weeks (n = 5 per strain per diet group). Hepatic retinoid concentrations were quantified by HPLC, and liver transcriptomes were profiled by bulk RNA sequencing. (B) Serum and hepatic retinoid concentrations in C57BL/6J and DBA/2J mice fed diets containing 5 or 25 IU vitamin A/g for 10 weeks. Dietary vitamin A supplementation increased hepatic RE levels without altering circulating ROH concentrations. (C) Principal component analysis of liver transcriptomes from C57BL/6J and DBA/2J mice fed diets containing 5 or 25 IU vitamin A/g. Samples clustered primarily according to strain, with PC1 accounting for 62% of total gene expression variance. (D) Permutation analysis of module gene expression responses to dietary vitamin A supplementation. Both the metSMAD and CYP3A modules showed significant but modest shifts in log₂ fold change distributions. (E) Expression responses of key retinoid-associated genes to dietary vitamin A supplementation. *Cyp26a1* was significantly induced by the 25 IU vitamin A/g diet, whereas the principal hub genes of the metSMAD and CYP3A modules, *Smad4* and *Cyp3a25*, respectively, remained largely unchanged in both strains.

As expected, the 25 IU vitamin A/g diet significantly increased hepatic RE levels in both strains, whereas serum ROH concentrations remained unchanged, reflecting the tight homeostatic regulation of circulating retinoids (**Fig. 6B**), regardless of the genotype. Consistent with the chow diet study (**Fig. 1B**; Tukey’s HSD-adjusted P = 1.0), hepatic RE levels were not significantly different between C57BL/6J and DBA/2J mice fed the same diet, either at 5 IU vitamin A/g (P = 0.662) or 25 IU vitamin A/g (P = 0.222). Despite substantial differences in hepatic RE concentrations between dietary groups, global liver transcriptional profiles were largely preserved, including the expression of retinoic acid receptor genes (**Fig. S16**). Consistent with the results from the strain comparison study (**Fig. 2A**), samples clustered primarily by strain, with PC1 accounting for 62% of total gene expression variance (**Fig. 6C**).

Permutation analysis revealed significant shifts in the distribution of log₂ fold changes among genes in both the metSMAD and CYP3A modules; however, the magnitude of these changes was modest (**Fig. 6D**). Moreover, although *Cyp26a1*, a canonical marker of elevated hepatic retinoic acid levels, was induced by the 25 IU vitamin A/g diet as expected, the principal hub genes of the two modules, *Smad4* (metSMAD module) and *Cyp3a25* (CYP3A module), showed little or no response to dietary vitamin A supplementation in either strain (**Fig. 6E**).

Together, these findings indicate that the transcriptional programs associated with genetically determined hepatic retinoid status are largely distinct from those induced by dietary vitamin A perturbation, suggesting that genetic background exerts a stronger influence on the metSMAD and CYP3A modules than moderate variation in dietary vitamin A intake.

### The CYP3A retinoid-associated gene module is selectively associated with pathological liver states

In addition to genetic and dietary influences, pathological liver conditions are known to alter hepatic retinoid homeostasis (reviewed by Melis et al.^55^). We therefore examined whether the metSMAD and CYP3A gene modules identified above are associated with transcriptional signatures of liver disease. We evaluated the relationship between liver disease phenotypes and expression of metSMAD and CYP3A module genes using a publicly available liver RNA-seq dataset generated from Diversity Outbred (DO) mice (GSE72759)^43,44^. Because DO mice capture extensive genetic variation, they offer an opportunity to assess disease-associated transcriptional responses while reducing strain-specific effects. The GSE72759 dataset includes liver gene expression profiles from 189 mice maintained on either a standard diet (SD, LabDiet 5K52 [protein: 20.1%cal, fat: 16.3%cal, carbohydrate: 61.6%cal, 20.0 IU vitamin A/g]) or a high-fat diet (HFD, Tekland TD.08811[protein: 14.7%cal, fat: 44.6%cal, carbohydrate: 40.7%cal, 7.6 IU vitamin A/g]): 49 female SD, 47 female HFD, 47 male SD, and 46 male HFD animals. We evaluated the contributions of reported covariates to transcriptional variation by regressing the first 10 principal components (PCs) derived from three expression datasets analyzed separately: all genes (top), CYP3A module hub genes (middle), and metSMAD module hub genes (bottom) (**Fig. S17**). The reported redox module eigengene showed significant associations in all analyses. Sex contributed significantly to variation in the expression of both the complete gene set and the CYP3A module hub genes, whereas dietary effects were detected only among the CYP3A module hub genes (**Fig. S17**). Nevertheless, the principal hub genes of the metSMAD and CYP3A modules, Smad4 and Cyp3a25, respectively, were consistently downregulated in HFD-fed mice regardless of sex (**Fig. S18AB**). Since *Cyp26a1*, a well-established marker of hepatic retinoic acid status, was also downregulated in HFD-fed mice (**Fig. S18C**), these findings suggest that HFD feeding may be associated with reduced hepatic retinoid signaling. Furthermore, the selective responsiveness of CYP3A module genes to HFD-induced transcriptional changes suggests that this module is sensitive to pathological alterations in liver physiology. This pattern was consistently observed in two independent liver transcriptomic datasets from patients with NAFLD (MAFLD), GSE193080^46,47^ and GSE268273^45^. In GSE193080, CYP3A module hub gene expression was significantly associated with the reported disease risk score (**Fig. S19**), whereas in GSE268273, CYP3A module hub genes were significantly associated with fibrosis stage and severity (**Fig. S20**). In contrast, metSMAD-module genes showed no significant associations in either dataset. Consistent with these results, the principal hub gene of the CYP3A module, *Cyp3a25*, was downregulated in both datasets (**Fig. S21**), whereas the principal metSMAD-module hub gene, *Smad4*, showed reduced expression only in GSE268273. These findings demonstrate that transcriptional changes in the CYP3A module consistently track liver disease severity across mouse and human datasets, whereas the metSMAD module remains comparatively stable. These results support the CYP3A module as a transcriptional signature associated with pathological liver states and altered hepatic retinoid homeostasis.

### Genetic background determines hepatic retinoid storage and circulating retinoid levels in mice by modulating mature liver cellular composition

Lastly, to investigate cell type-specific transcriptional differences associated with hepatic retinoid status, we performed single-nucleus RNA-seq (snRNA-seq) using the 10x Genomics platform on three wild-derived mouse strains, CAST/EiJ, WSB/EiJ, and PWK/PhJ, representing strains with the highest and lowest hepatic RE levels. Based on their transcriptional profiles, 13 cell clusters were identified, including five hepatocyte subtypes and two hepatic stellate cell (HSC) subtypes (**Fig. 7AB; Fig. S22; Fig. S23**). As expected, each strain exhibited a distinct cellular composition in the adult liver (**Fig. 7C**).

**Fig. 7.**
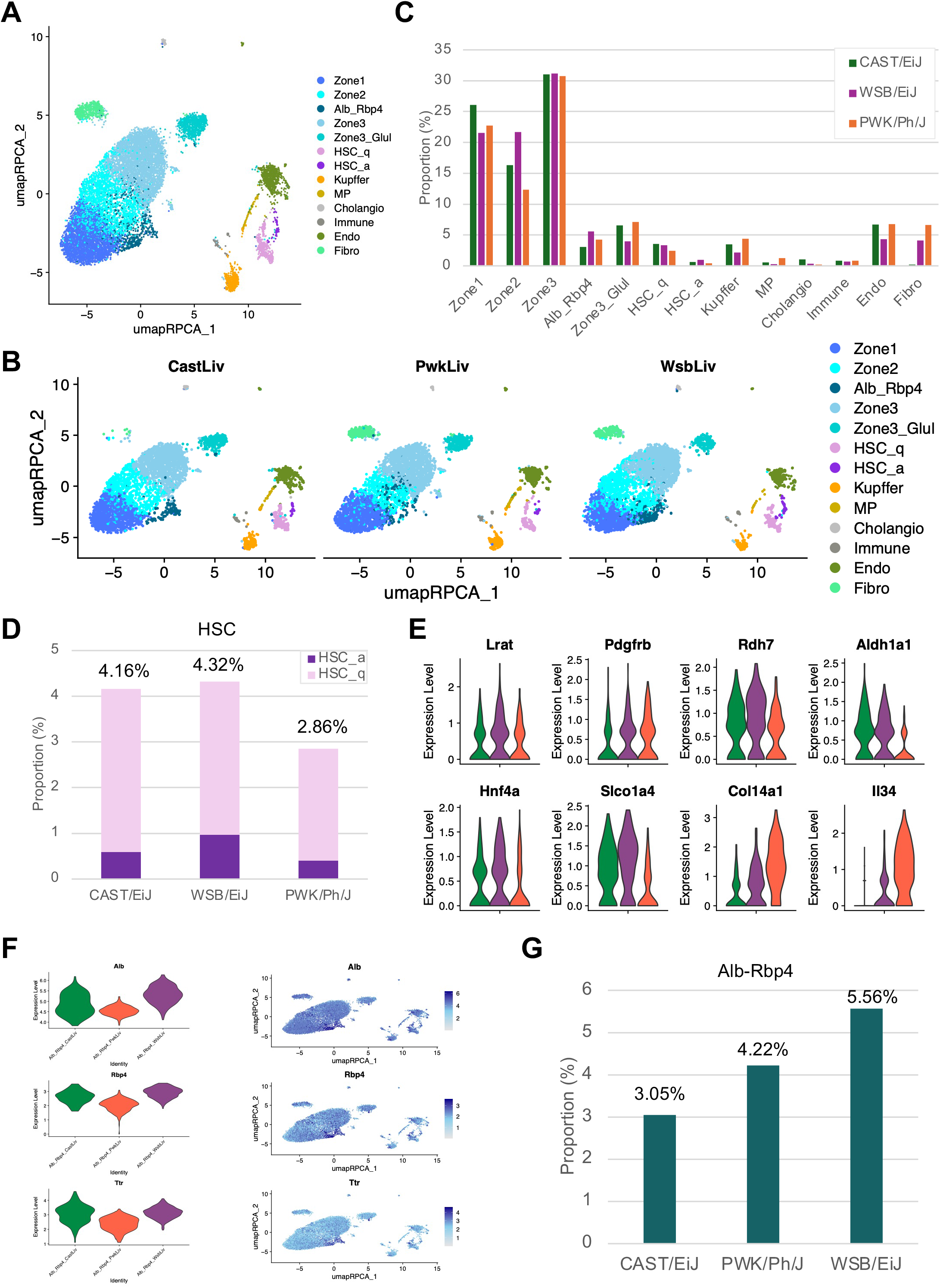
Genetic background is associated with differences in liver cellular composition and retinoid-related cell populations. (A) UMAP visualization of single-nucleus RNA-seq profiles from CAST/EiJ, WSB/EiJ, and PWK/PhJ livers. Thirteen transcriptionally distinct cell populations were identified, including five hepatocyte subtypes and two hepatic stellate cell (HSC) subtypes. (B) UMAP plots showing cells from each strain separately (CAST/EiJ, WSB/EiJ, and PWK/PhJ), illustrating strain-dependent differences in the representation of liver cell populations. (C) Relative abundance of liver cell populations in CAST/EiJ, WSB/EiJ, and PWK/PhJ mice determined by single-nucleus RNA sequencing. Distinct cellular compositions were observed among strains, indicating substantial genetic influences on adult liver cell population structure. (D) Proportion of hepatic stellate cells (HSCs) in CAST/EiJ, WSB/EiJ, and PWK/PhJ livers. PWK/PhJ mice exhibited a lower overall abundance of HSCs than CAST/EiJ and WSB/EiJ mice despite similar proportions of activated and quiescent HSC subtypes. (E) Expression of selected retinoid-related genes (*Lrat*, *Rdh7*, *Aldh1a1*), hepatic stellate cell marker genes (*Pdgfrb*), hepatocyte-associated genes (*Hnf4a* and *Slco1a4*), and fibrosis-associated genes (*Col14a1* and *Il34*) in hepatic stellate cells from CAST/EiJ, WSB/EiJ, and PWK/PhJ mice. (F) Expression of *Alb*, *Rbp4*, and *Ttr* in the Alb-Rbp4 hepatocyte population identified by snRNA-seq. (G) Association between Alb-Rbp4 hepatocyte abundance and serum retinol (ROH) concentrations across strains.

Intriguingly, although the proportions of activated and quiescent HSCs were comparable among strains, the overall abundance of HSCs was lower in PWK/PhJ (2.86%), the strain with the lowest hepatic RE levels, than in CAST/EiJ (4.16%) and WSB/EiJ (4.32%), both of which exhibit higher hepatic RE storage (**Fig. 7D**). This difference was independently confirmed by cell deconvolution analysis of bulk liver RNA-seq data using CIBERSORTx ^56^(**Fig. S24**). Differential expression analysis of HSCs further showed that *Lrat* expression, which encodes the main enzyme responsible for retinol esterification and RE synthesis, did not differ substantially among strains (**Fig. 7E**). These findings suggest that the reduced hepatic RE storage capacity of PWK/PhJ is unlikely to result from differences in *Lrat* expression and instead may be attributable, at least in part, to reduced HSC abundance. Rather, it may reflect the reduced abundance of HSCs, the primary cellular reservoir for hepatic RE storage.

Although no histopathological evidence of fibrosis was detected (**Fig. S25**), *Col14a1* and *Il34*, genes implicated in fibrogenic processes, were significantly upregulated in PWK/PhJ HSCs (**Fig. 7E**). Consistent with this observation, genes more highly expressed in PWK/PhJ were enriched for pathways involved in cytoskeletal organization, integrin signaling, focal adhesion, and vascular smooth muscle contraction, whereas genes upregulated in CAST/EiJ and WSB/EiJ were enriched predominantly for metabolic pathways (**Fig. S26**). Together, these findings suggest that PWK/PhJ mice exhibit transcriptional signatures associated with fibrogenic remodeling despite the absence of overt histological evidence of fibrosis. These molecular features may contribute to the previously reported susceptibility of this strain to metabolic-associated steatohepatitis (MASH)^57^.

Among hepatocyte populations, one cluster characterized by high *Alb* expression, the Alb-Rbp4 cluster, also showed elevated *Rbp4* and *Ttr* expression (**Fig. 7F**), two genes that play central roles in systemic retinoid circulation. Notably, the abundance of Alb-Rbp4 cells was associated with serum ROH levels (**Fig. 7G**). Furthermore, 10 of the top 20 serum ROH-associated genes identified in the bulk RNA-seq analysis, including 6 of the top 10 genes, were Alb-Rbp4 cluster marker genes (**Table S10**). These findings suggest that this hepatocyte subpopulation contributes disproportionately to regulating circulating retinoid levels.

Taken together, these findings indicate that genetic background influences hepatic retinoid storage and circulating retinoid levels, at least in part through differences in mature liver cellular composition. Specifically, variation in HSC abundance is associated with hepatic RE storage, whereas variation in an Rbp4/Ttr-enriched hepatocyte population is associated with circulating ROH levels.

## Discussion

We demonstrated that genetic background influences retinoid storage and circulation and is associated with distinct molecular programs and differences in adult liver cellular composition. While previous human family studies have reported heritable effects on circulating retinoid concentrations^13,15^, and a mouse study using a limited panel of inbred strains suggested strain-dependent variation in vitamin A status^25^, to our knowledge, this is the first study to comprehensively characterize genetic influences on both circulating retinoid levels and hepatic retinoid storage in mice.

This study also attempted to corroborate these findings in liver transcriptomic datasets from patients with liver diseases. These observations underscore the importance of integrating genetic information into precision nutrition approaches to assess and manage vitamin A status.

Single-nucleus RNA-seq analysis of three wild-derived mouse strains demonstrated that liver cellular composition varies substantially by genetic background. The proportion of HSCs, the primary storage site for hepatic retinoids, was positively associated with hepatic RE levels, whereas *Lrat* expression in HSCs did not differ appreciably among strains. Because these analyses were limited to three wild-derived strains, broader genetic differences among subspecies may contribute to the observed variation. Nevertheless, the positive association between HSC abundance and hepatic RE levels suggests that cellular composition may be an important determinant of retinoid storage capacity. In addition, the proportion of a hepatocyte subpopulation with high expression of *Rbp4* and *Ttr*, two key serum retinol transport proteins ^58^, was positively associated with circulating retinoid levels. These results indicate that genetic variation may influence retinoid storage and circulation by shaping the cellular architecture of the liver. Interestingly, the association between hepatic RE levels and HSC abundance aligns with the relationship between hepatic RE storage and the metSMAD module, suggesting that genetic differences in liver cellular composition and transcriptional regulation may jointly contribute to variation in retinoid storage.

Our WGCNA analysis identified two gene modules associated with genetically determined hepatic retinoid status. The metSMAD module was associated with liver metabolic profiles and enriched for genes downstream of *Smad4*, whereas the CYP3A module showed no association with liver metabolic profiles. Among the CYP3A module hub genes, *Cyp3a25* showed a strong association with *Rarb* and *Rarg* expression. *Cyp3a25* is a homolog of human *CYP3A4*, together with *Cyp3a11*^59^. Although *Cyp3a11* is expressed at substantially higher levels than *Cyp3a25*, its expression was not associated with hepatic RE levels. Both *Cyp3a25* and *Cyp3a11* are regulated by the pregnane X receptor (PXR)^60^. PXR is a key regulator of genes involved in xenobiotic metabolism, including *CYP3A4*, as well as pathways related to energy homeostasis, inflammation, and bile acid metabolism^61^. These findings suggest that genes within the CYP3A module may reflect alterations in hepatic metabolic and regulatory processes associated with liver physiological or pathological status. The metSMAD module was enriched for SMAD4 target genes, and hepatic Smad4 expression was positively associated with RE concentrations, identifying SMAD4 as a candidate upstream regulator of the transcriptional network associated with retinoid storage. However, this study was not designed to directly test developmental mechanisms, and future studies will be needed to determine whether SMAD4 causally contributes to differences in hepatic retinoid storage.

Notably, expression of genes in neither module was substantially altered by dietary vitamin A supplementation. In contrast, pathological liver conditions, including steatosis and fibrosis, selectively affected expression of CYP3A module genes, whereas metSMAD-module genes remained largely unchanged. Together, these observations indicate that the metSMAD and CYP3A modules capture distinct aspects of hepatic retinoid biology. The metSMAD module appears to reflect genetically determined variation in hepatic retinoid storage and liver metabolism, whereas the CYP3A module is more responsive to pathological perturbations and consistently tracks disease-associated alterations in hepatic retinoid homeostasis.

Excessive accumulation of vitamin A in the liver is known to cause hepatocyte injury, necrosis, and activation and hyperplasia of the primary storage cells for RE, hepatic stellate cells (HSCs), leading to fibrosis^62^. Approximately 70-80% of retinoids in the body are stored in the liver, and of those, 90-95% are stored in lipid droplets within HSCs^63^. Pathologically, vitamin A toxicity progresses from hepatic steatosis to fibrosis and cirrhosis^62^. Intriguingly, among the strains we tested, PWK/Ph/J showed the lowest liver retinoid storage level, and CAST/EiJ has the highest, with WSB/EiJ being intermediate. Notably, these strains are known to have different susceptibilities to metabolic defects caused by high-fat or Western diets (reviewed by Masson et al.^57^). PWK/Ph/J is prone to metabolic defects induced by such diets, particularly metabolic-associated steatohepatitis (MASH). In contrast, WSB/EiJ and CAST/EiJ are resistant to these metabolic conditions. These observations raise the possibility that genetically determined differences in physiological hepatic retinoid storage contribute to variation in susceptibility to metabolic liver disease.

This study has several limitations. First, retinoid measurements and molecular assays were performed exclusively in female mice. Previous studies have reported sexual dimorphism in circulating retinoid-binding protein levels and sex-dependent differences in retinoid storage and metabolism during development^64,65^. Inclusion of male mice in future studies may provide a more comprehensive understanding of the genetic determinants of retinoid homeostasis. Second, although we identified significant genetic influences on circulating and hepatic retinoid levels and uncovered candidate regulatory factors associated with these traits, using inbred mouse strains limited our ability to map the specific genetic loci responsible for the observed variation. Future studies using genetically diverse outbred populations, such as Diversity Outbred mice, will be necessary to identify causal genomic regions and genes underlying retinoid-related phenotypes. Although several established HSC and HSC-enriched marker genes identified in single-cell RNA-seq studies, including known quiescent markers *Gfap*, *Ntm* and *Ngfr*^66^, *Col3a1*^67^, and *Hhip*^68^, showed positive associations with hepatic RE or ROH levels in our bulk RNA-seq analysis of 20 inbred strains, the single-nucleus RNA-seq analysis was restricted to three wild-derived strains selected for their extreme hepatic RE phenotypes. Consequently, the extent to which these cellular findings generalize across the broader strain panel remains uncertain. Therefore, the observed relationships between liver cellular composition and retinoid status should be validated in a broader panel of genetically diverse strains. Third, although our dietary study found that the genetically associated modules differ from diet-responsive modules, it tested only two common lab inbred strains, and the sample size is relatively small. Therefore, we cannot fully rule out the possibility that our findings are limited to those two strains or that we missed subtle alterations due to limited statistical power. Finally, although we replicated the disease-associated transcriptional alterations of the CYP3A module in two independent human liver transcriptomic datasets, the extent to which our findings can be generalized to human populations remains uncertain, as the experimental approaches used in this study are not readily feasible in humans. Additional studies in human cohorts are needed to validate these molecular signatures and determine their relevance to human retinoid biology, liver disease progression, and precision nutrition.

In summary, although dietary vitamin A intake is the primary determinant of retinoid status, genetic variation substantially contributes to individual differences in circulating retinoid concentrations and hepatic retinoid storage, highlighting genetics’ important role in regulating retinoid homeostasis. We identify distinct transcriptional programs associated with hepatic retinoid storage and disease-associated retinoid dysregulation and provide evidence that variation in mature liver cellular composition contributes to both hepatic and circulating retinoid phenotypes. These findings expand our understanding of retinoid biology beyond dietary exposure alone and suggest that genetically determined differences in hepatic retinoid homeostasis may contribute to inter-individual variation in susceptibility to metabolic liver disease. Incorporating genetic information into assessments of vitamin A status may therefore be an important step toward the development of precision nutrition strategies tailored to individual metabolic and physiological characteristics.

## Data availability

The raw FASTQ files and processed sequencing data will be available via the Gene Expression Omnibus (GEO). Untargeted metabolomics data are available upon request.

## Code availability

All analyses were conducted using publicly available software. The code used to perform the analysis from the raw data is available upon request.

## Acknowledgments

We thank the Institute of Animal Studies at Albert Einstein College of Medicine for their animal husbandry services. Additionally, portions of this research were conducted using the advanced computing resources provided by Texas A&M High Performance Research Computing. The contributions of Dr. Yuta Matsuno to this study were made while he was affiliated with Texas A&M University. His current affiliation is the Department of Animal Science, Faculty of Agriculture, Tokai University, Kumamoto, Japan.

## Declaration of Generative AI and AI-assisted technologies in the writing process

During the preparation of this work, we used Microsoft Copilot, Grammarly, and DeepL in order to improve grammar and English expression. After using this tool/service, we reviewed and edited the content as needed and took full responsibility for the content of the publication.

## Author contributions

M.S. conceived and led the study. W.V.C., L.Q., D.W.T., and M.S. designed the research framework. Y.M., M.A.G., and M.S. performed animal treatment and production. A.K.S.K., and M.S. performed integrative transcriptomic and metabolomic analyses. A.S. and Y.S. performed validation studies. Y.K. and L.Q. performed retinoid measurements. Y.M., A.K.S.K., Y.K., A.S., Y.S., B.G., W.V.C., L.Q., D.W.T., and M.S. interpreted the data. Y.M., A.K.S.K., and M.S. drafted the paper. W.V.C., L.Q., and D.W.T. provided overall supervision. All authors critically reviewed and approved the final paper.

## List of Abbreviations

AIN-93: American Institute of Nutrition 1993 rodent diet formulation
ALP: Alkaline phosphatase
ALT: Alanine aminotransferase
ANOVA: Analysis of variance
ARRIVE: Animal Research: Reporting of In Vivo Experiments
AST: Aspartate aminotransferase
CC: Collaborative Cross
CIBERSORTx: Cell-type Identification By Estimating Relative Subsets Of RNA Transcripts
DESeq2: Differential Expression Sequencing 2
DO: Diversity Outbred
eQTL: Expression quantitative trait locus
FDR: False discovery rate
FASTQ: FastQ sequence file format
GB: Gigabase(s)
GEO: Gene Expression Omnibus
GO: Gene Ontology
GPCR: G protein-coupled receptor
GTEx: Genotype-Tissue Expression
H²: Broad-sense heritability
HDL: High-density lipoprotein
HMDB: Human Metabolome Database
HPLC: High-performance liquid chromatography
HSC: Hepatic stellate cell
IACUC: Institutional Animal Care and Use Committee
KEGG: Kyoto Encyclopedia of Genes and Genomes
MASLD: Metabolic dysfunction-associated steatotic liver disease
MAST: Model-based Analysis of Single-cell Transcriptomics
MASH: Metabolic dysfunction-associated steatohepatitis
MM: Module membership
MPD: Mouse Phenome Database
MSEA: Metabolite set enrichment analysis
MS: Metabolite significance (Fig. 5B)
mTOR: Mechanistic target of rapamycin
NAFLD: Nonalcoholic fatty liver disease
NEFA: Non-esterified fatty acids
ORA: Overrepresentation analysis
PCA: Principal component analysis
PC: Principal component
PXR: Pregnane X receptor
qRT-PCR: Quantitative reverse transcription polymerase chain reaction
RA: Retinoic acid
RaMP-DB: Relational Database of Metabolomic Pathways
RAR: Retinoic acid receptor
RBP4: Retinol-binding protein 4
RE: Retinyl ester
ROH: Retinol
RNA-seq: RNA sequencing
RXR: Retinoid X receptor
SD: Standard diet
SNP: Single nucleotide polymorphism
snRNA-seq: Single-nucleus RNA sequencing
SRA: Sequence Read Archive
STRA6: Stimulated by retinoic acid 6
TF: Transcription factor
TTR: Transthyretin
UMAP: Uniform Manifold Approximation and Projection
UPLC-MS/MS: Ultra-performance liquid chromatography tandem mass spectrometry
WGCNA: Weighted Gene Correlation Network Analysis
WMCNA: Weighted Metabolite Correlation Network Analysis

---

**Fig. S1.**
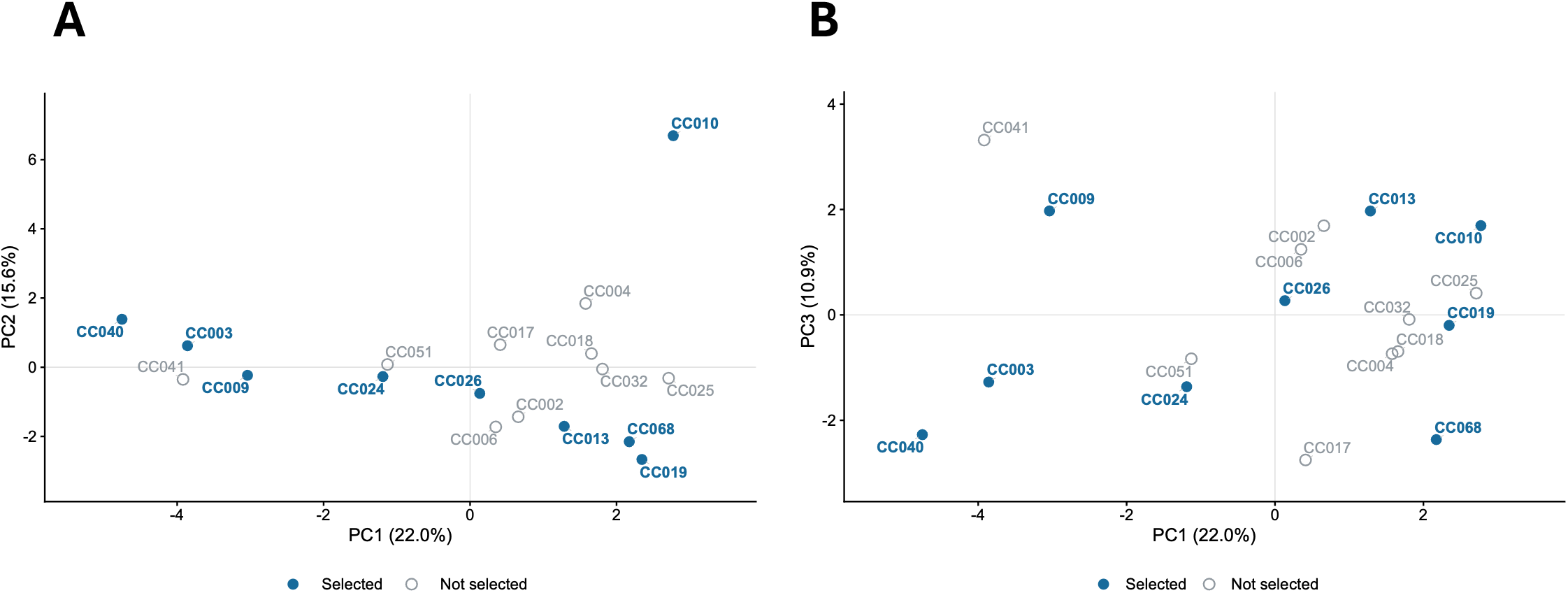
Phenotypic distribution of selected Collaborative Cross strains. Principal component analysis (PCA) of five body size and composition traits and 22 blood chemistry traits was performed using publicly available female strain means from 18 Collaborative Cross (CC) strains (JaxCC1, Mouse Phenome Database, MPD:519). All traits were centered and scaled to unit variance before PCA. (A) PC1 versus PC2 and (B) PC1 versus PC3. The nine CC strains selected for the present study are highlighted in blue, while the remaining nine strains are shown in gray. Percentages on the axes indicate the variance explained by each principal component.

**Fig. S2.**
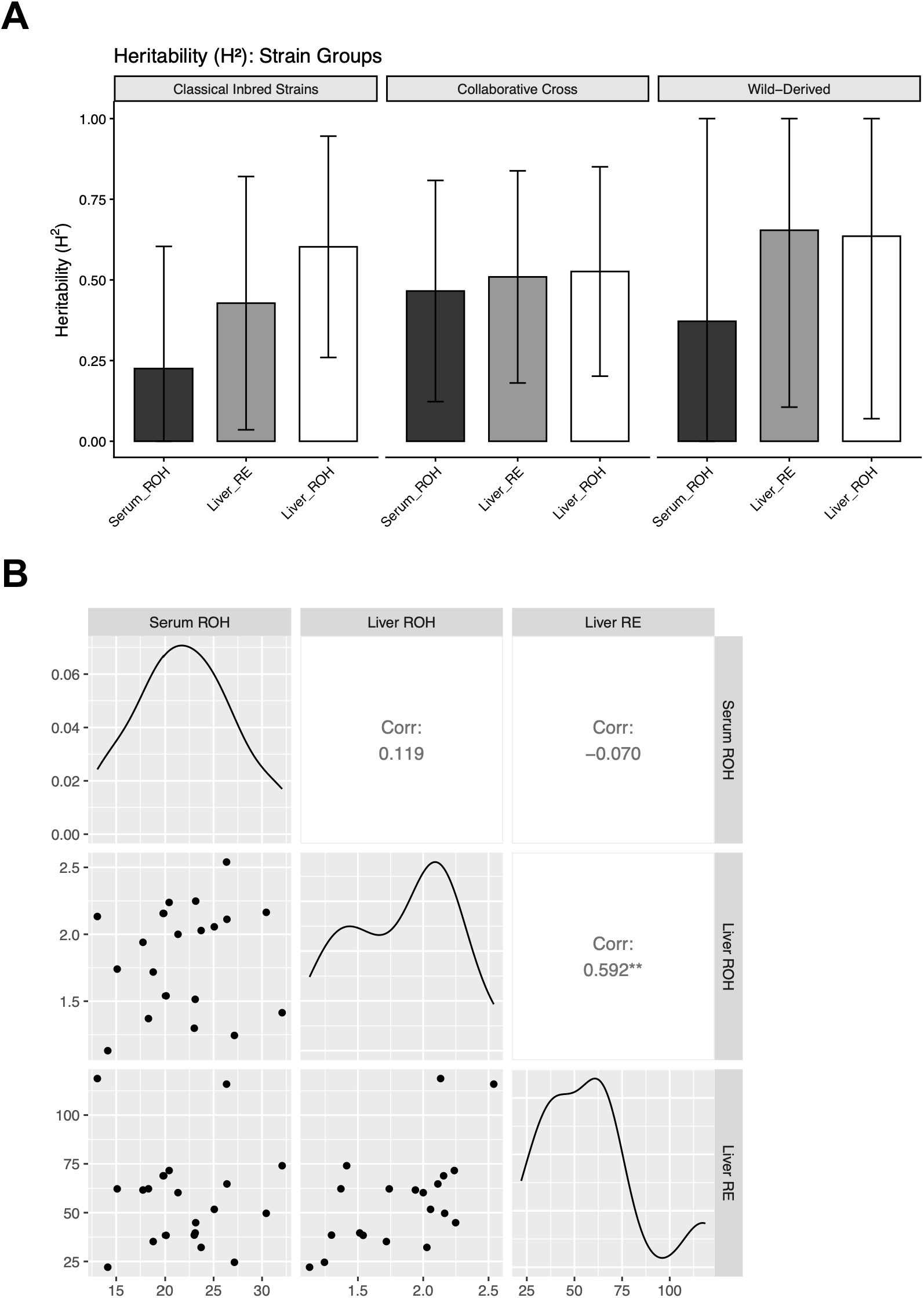
Heritability and correlation analyses of circulating and hepatic retinoid traits across mouse strain groups. **(A)** Comparison of broad-sense heritability (H²) estimates for serum retinol (ROH), hepatic retinol (ROH), and hepatic retinyl ester (RE) among classical laboratory, wild-derived, and Collaborative Cross (CC) mouse strains. No significant differences in heritability estimates were observed among strain groups. **(B)** Correlation analysis of strain-averaged serum retinol (ROH), hepatic retinol (ROH), and hepatic retinyl ester (RE) concentrations across the 20 inbred mouse strains. Consistent with the individual-animal analysis shown in Fig. 1D, hepatic ROH and RE concentrations were positively correlated, whereas serum ROH was not significantly correlated with either hepatic retinoid measure.

**Fig. S3.**
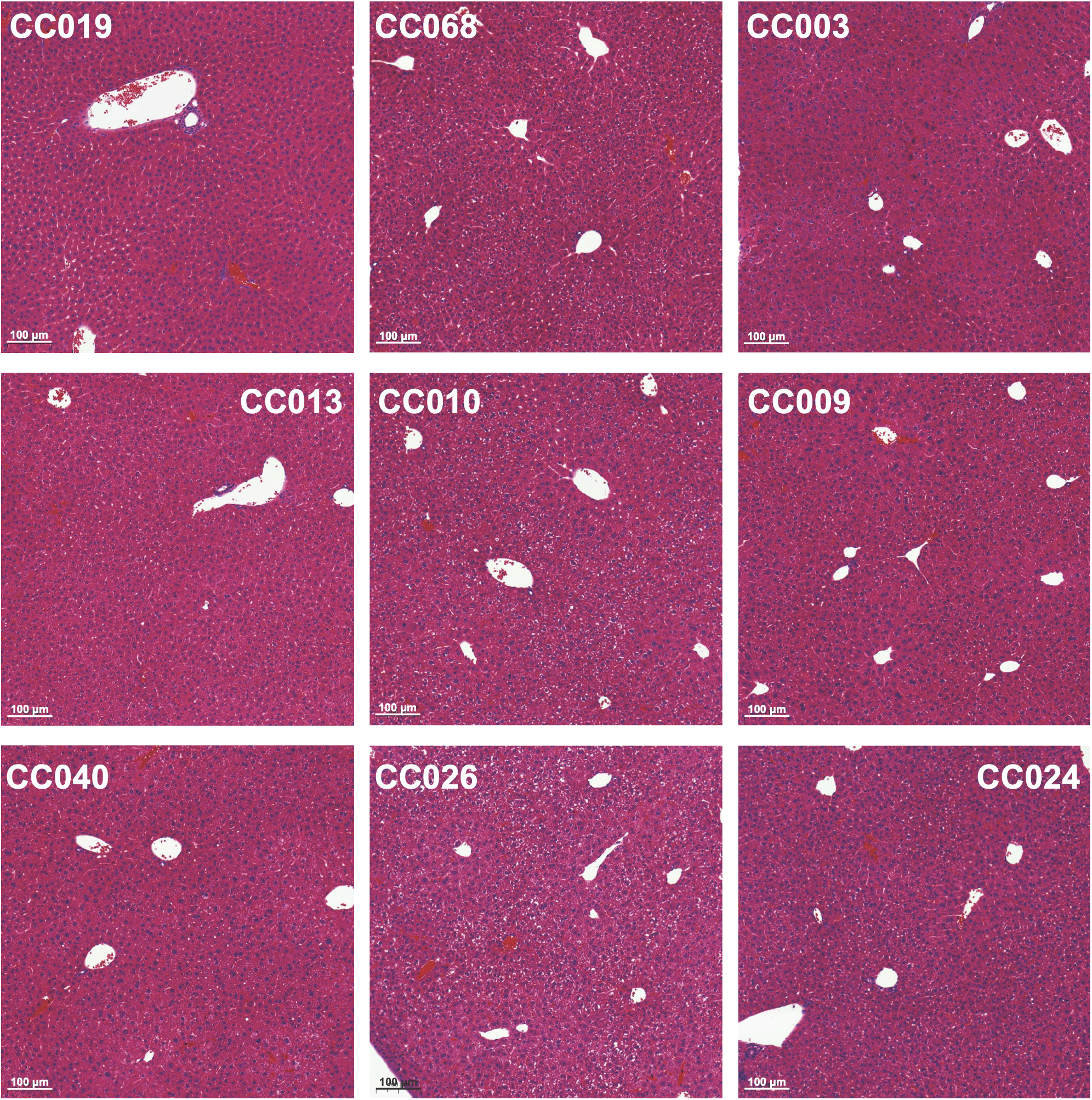
Histopathological assessment of livers from CC strains. A histopathological assessment was performed on liver samples from nine CC strains. Standard H&E staining was used for the assessment. Each panel shows a representative image of a strain. The panels are ordered from the highest to the lowest hepatic RE levels, from top left to bottom right. No difference associated with hepatic RE levels was observed.

**Fig. S4.**
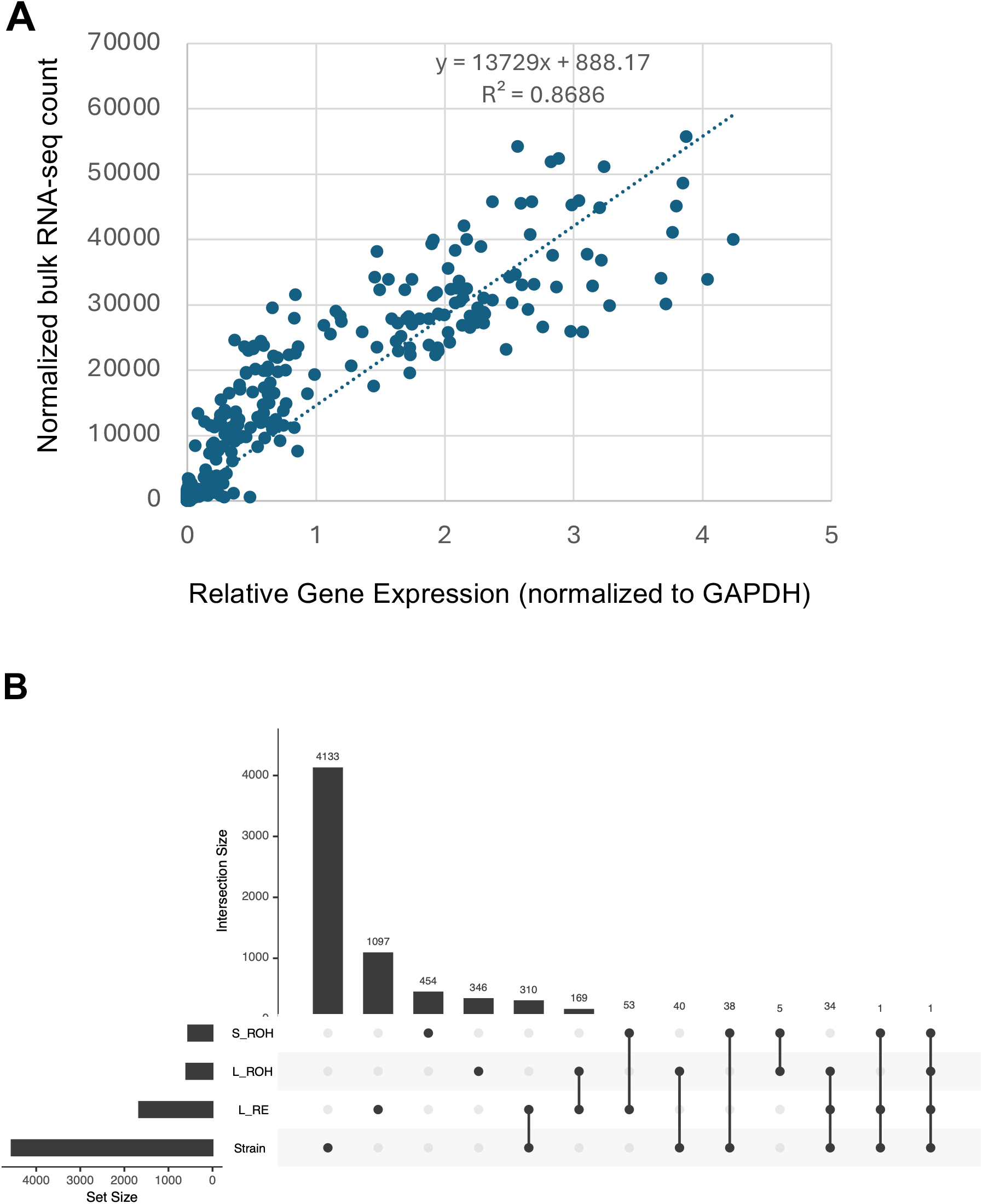
Validation of transcriptomic data and overlap analysis of strain-dependent and retinoid-associated genes. **(A)** Validation of RNA-seq gene expression measurements by quantitative RT-PCR (qRT-PCR). Relative gene expression levels normalized to *Gapdh* were strongly correlated with normalized RNA-seq counts across samples (R² = 0.8686), demonstrating high concordance between the two methods. **(B)** UpSet plot showing the overlap among genes exhibiting significant strain-dependent expression and genes significantly associated with serum retinol (ROH), hepatic retinol (ROH), or hepatic retinyl ester (RE) concentrations. Hepatic RE-associated genes showed the greatest overlap with strain-dependent genes (345 genes), followed by hepatic ROH-associated genes (74 genes) and serum ROH-associated genes (39 genes).

**Fig. S5.**
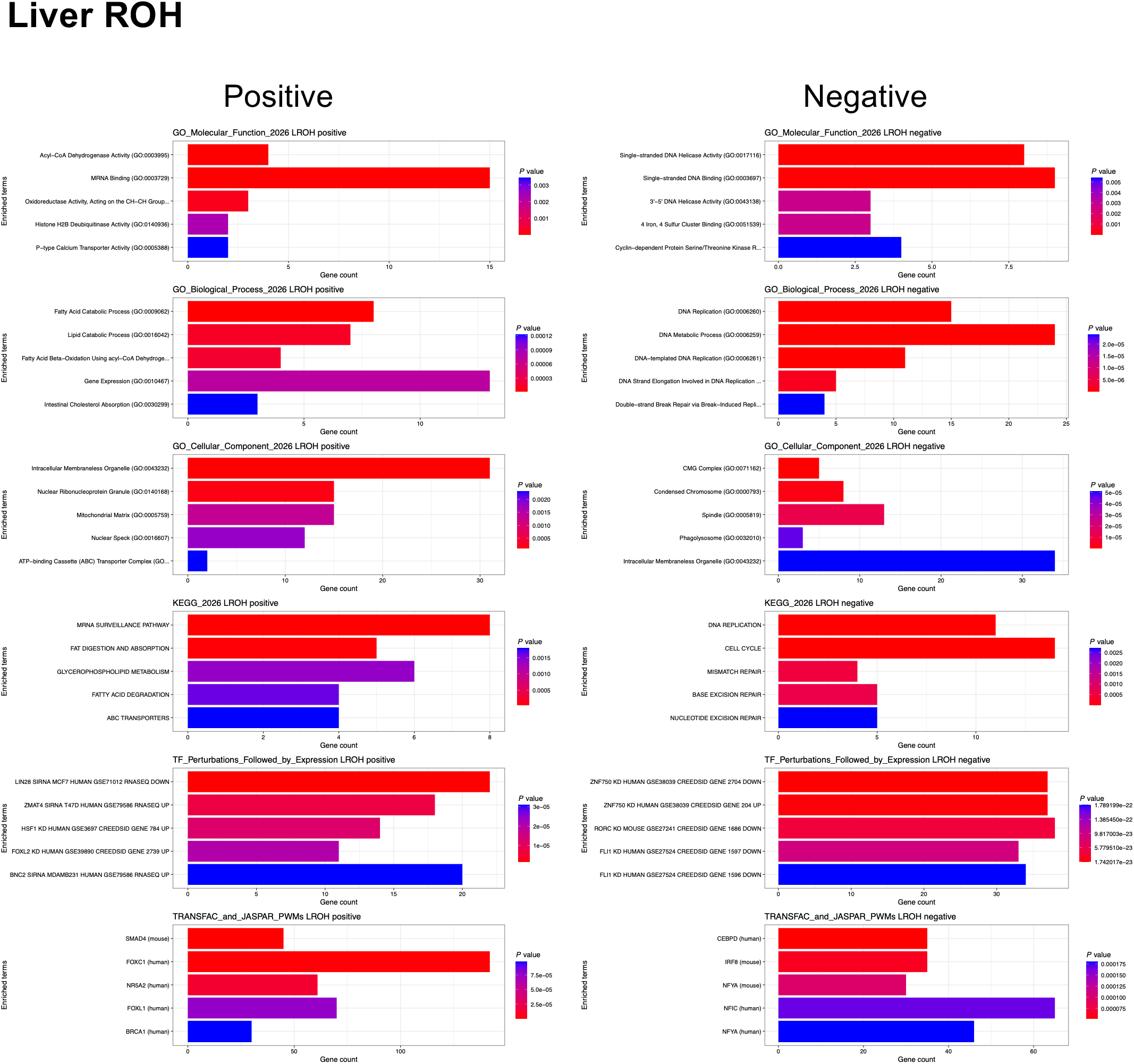
Functional enrichment analysis of liver retinol-associated genes. Left, enrichment analysis of genes positively associated with hepatic retinol (ROH) concentrations. Right, enrichment analysis of genes negatively associated with hepatic ROH concentrations. Rows show Gene Ontology (GO) molecular function, GO biological process, GO cellular component, KEGG pathways, transcription factor perturbation signatures, and TRANSFAC transcription factor motif enrichment analyses.

**Fig. S6.**
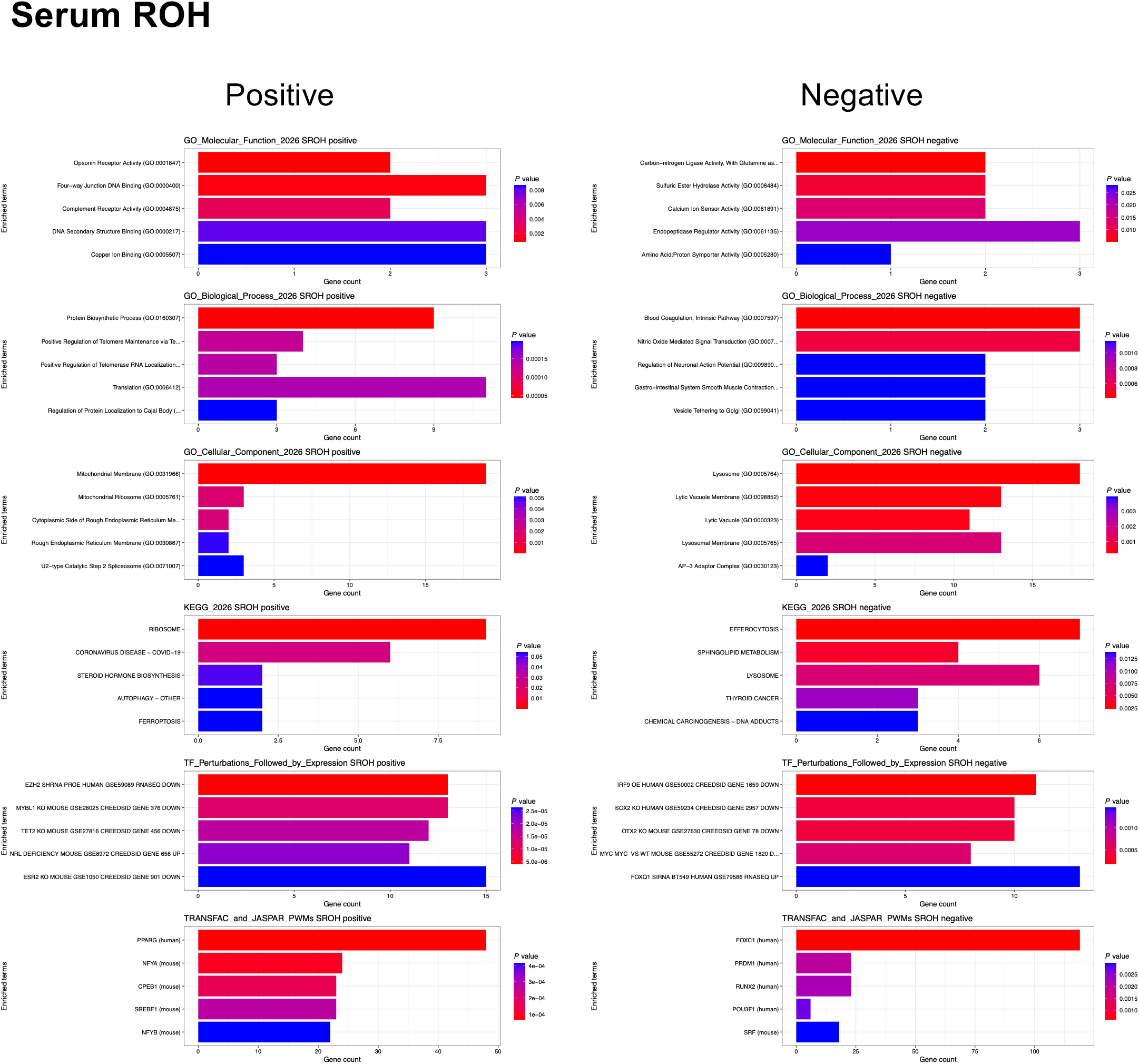
Functional enrichment analysis of serum retinol-associated genes. Left, enrichment analysis of genes positively associated with serum retinol (ROH) concentrations. Right, enrichment analysis of genes negatively associated with serum ROH concentrations. Rows show Gene Ontology (GO) molecular function, GO biological process, GO cellular component, KEGG pathways, transcription factor perturbation signatures, and TRANSFAC transcription factor motif enrichment analyses.

**Fig. S7.**
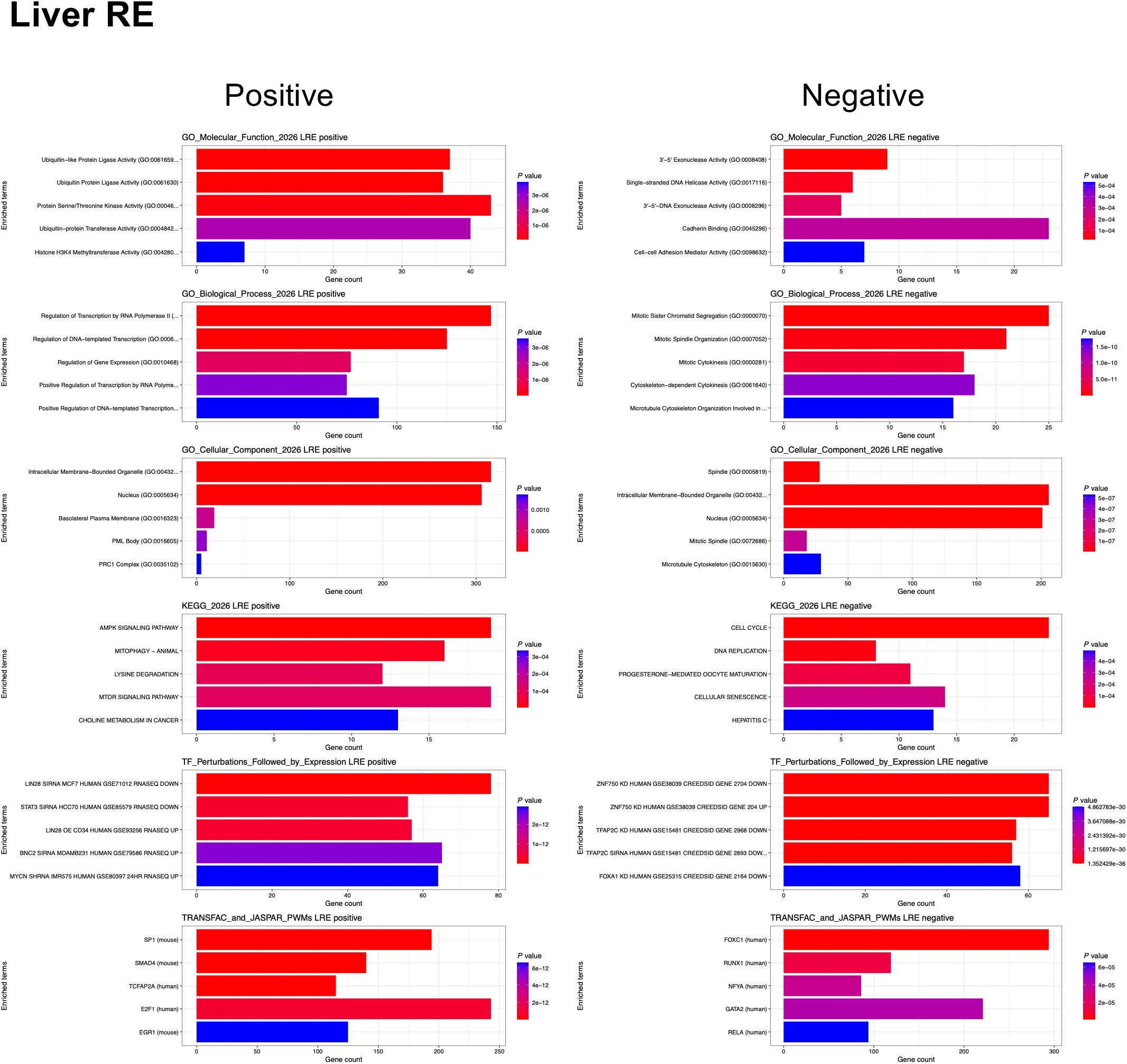
Functional enrichment analysis of hepatic retinyl ester-associated genes. Left, enrichment analysis of genes positively associated with hepatic retinyl ester (RE) concentrations. Right, enrichment analysis of genes negatively associated with hepatic RE concentrations. Rows show Gene Ontology (GO) molecular function, GO biological process, GO cellular component, KEGG pathways, transcription factor perturbation signatures, and TRANSFAC transcription factor motif enrichment analyses.

**Fig. S8.**
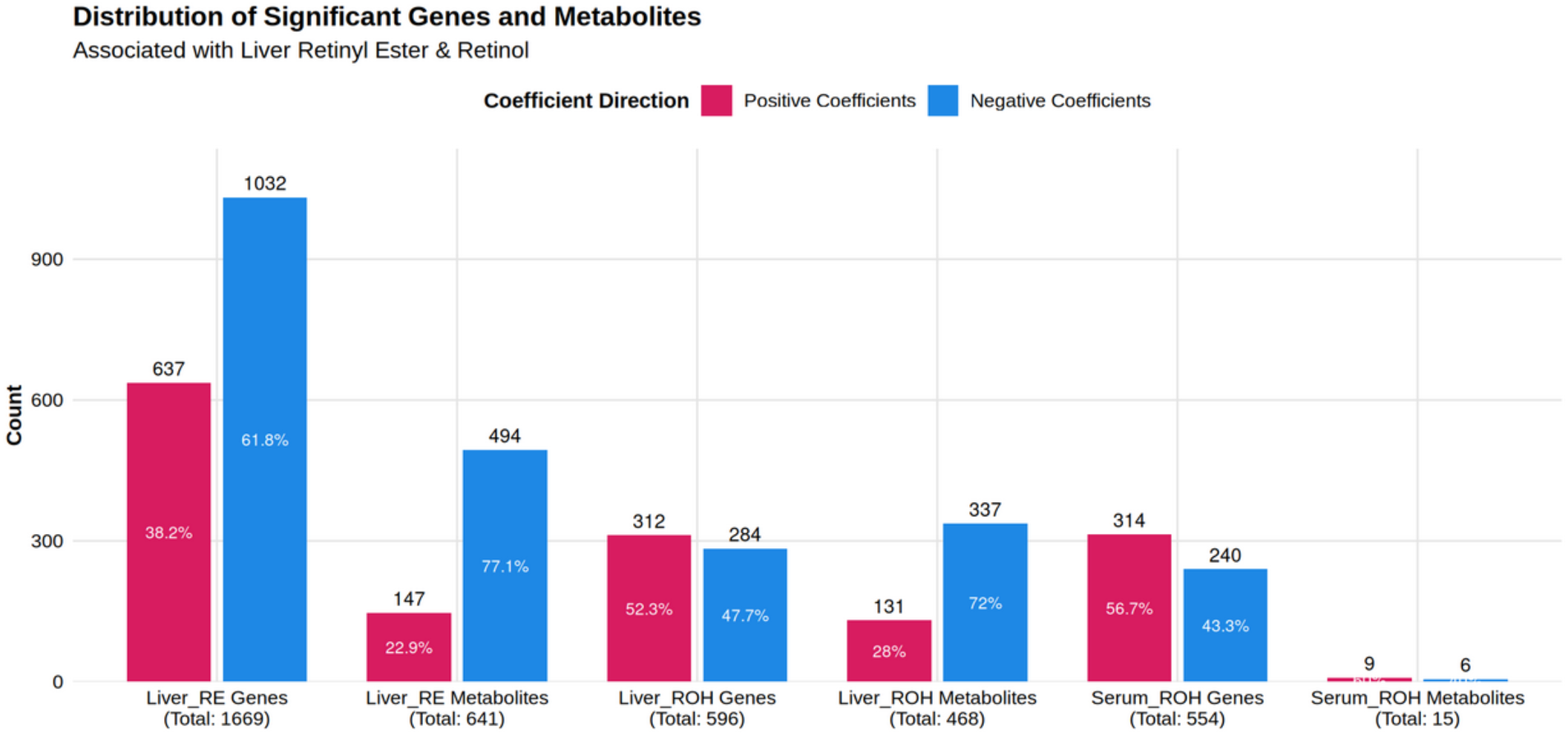
Transcriptomic associations with retinoid status exceed metabolomic associations. Comparison of the numbers of significantly associated genes and metabolites for serum retinol (ROH), hepatic retinol (ROH), and hepatic retinyl ester (RE) concentrations. For all retinoid measures, substantially more genes than metabolites were significantly associated with retinoid status.

**Fig. S9.**
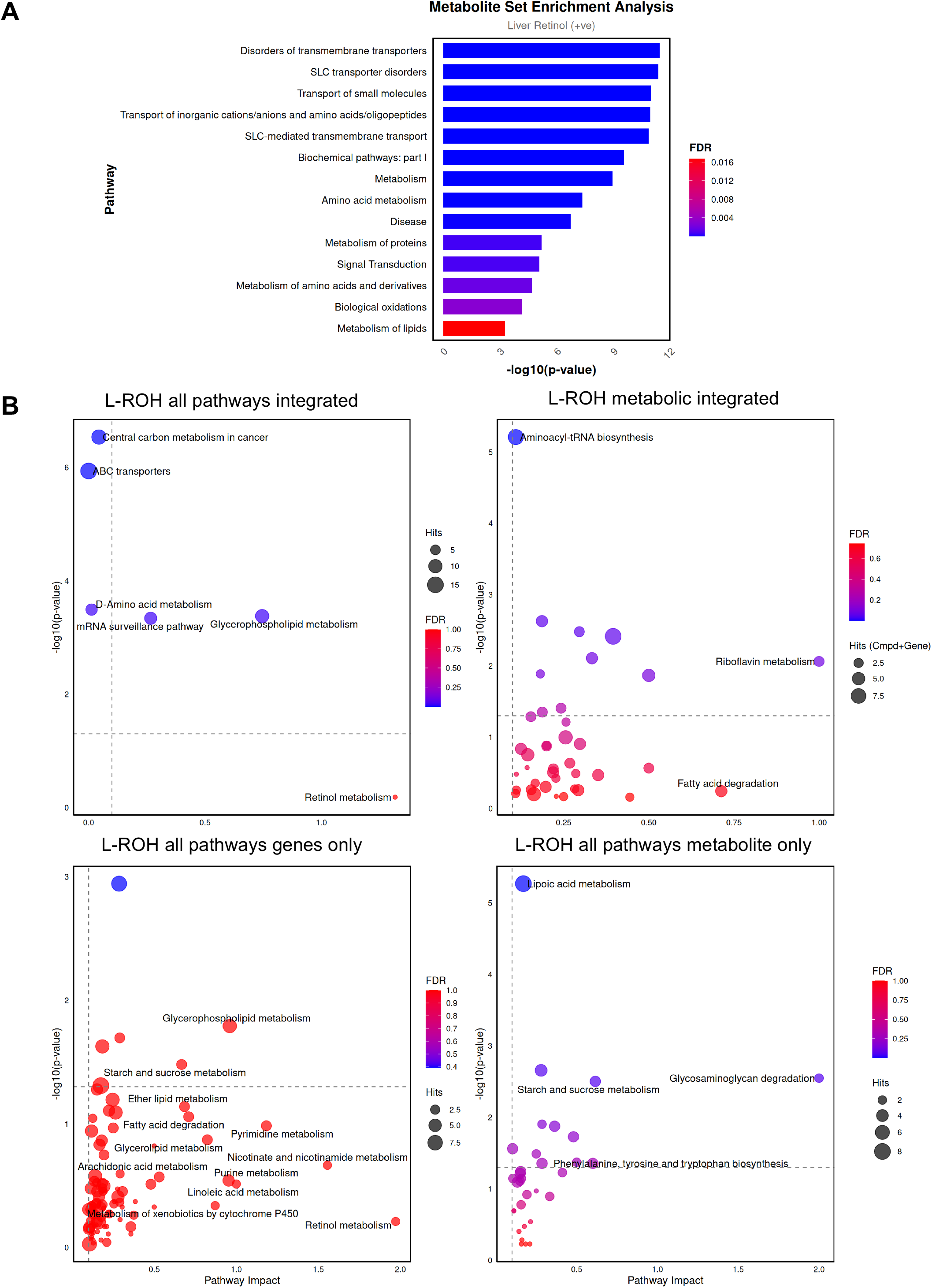
Pathway analyses identify molecular networks associated with hepatic retinol concentrations. **(A)** Metabolite set enrichment analysis of metabolites positively associated with hepatic retinol (ROH) concentrations. ROH-associated metabolites were enriched in transporter-and amino acid metabolism-related pathways. **(B)** Integrated pathway analysis of hepatic ROH-associated transcriptomic and metabolomic data. Top left, significantly enriched pathways associated with hepatic ROH levels. Top right, pathways jointly supported by both gene and metabolite datasets. Bottom left, pathways supported exclusively by transcriptomic data. Bottom right, pathways supported exclusively by metabolomic data.

**Fig. S10.**
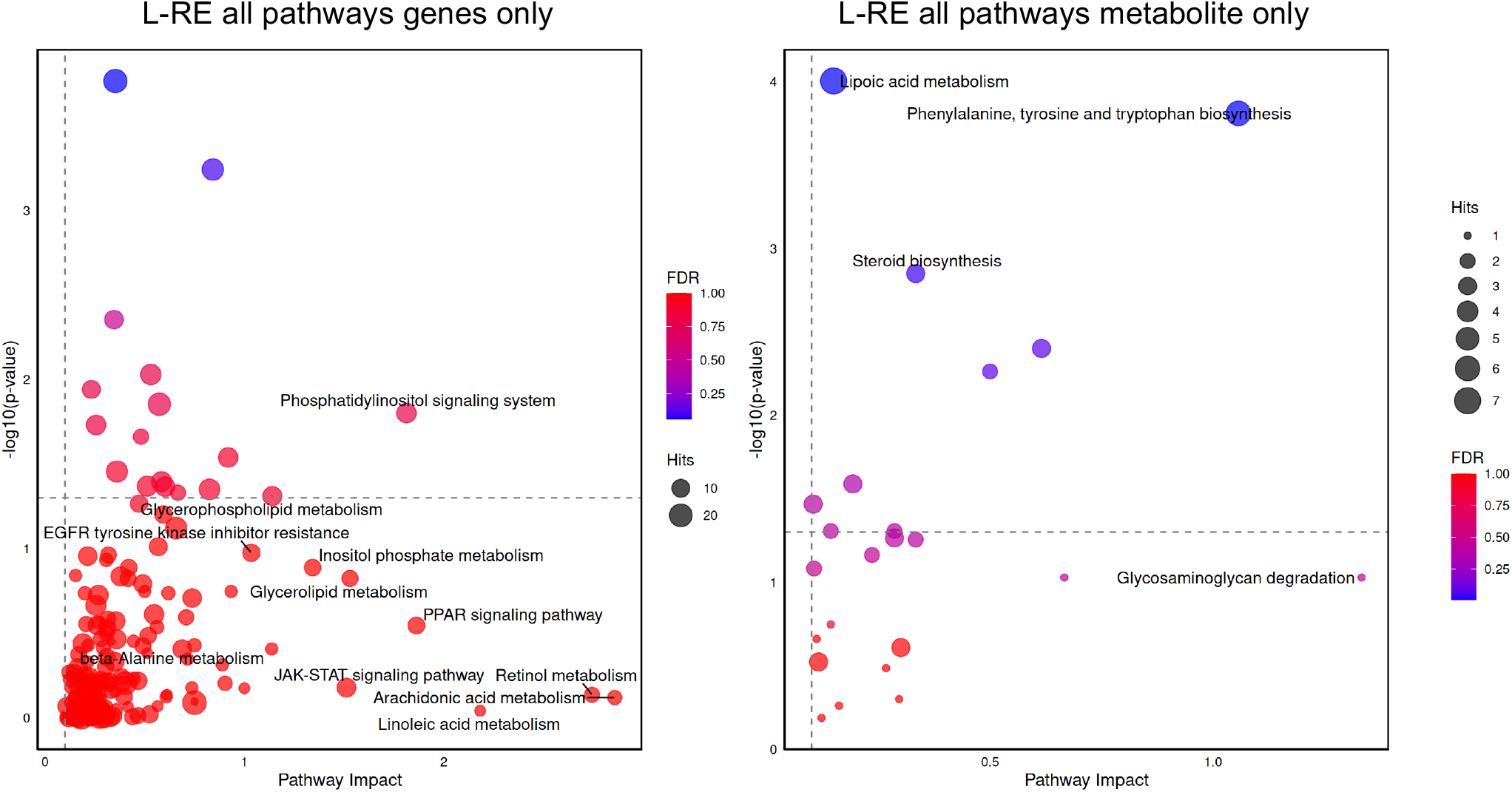
Transcriptome-specific and metabolome-specific pathways associated with hepatic retinyl ester concentrations. Integrated pathway analysis of hepatic retinyl ester (RE)-associated transcriptomic and metabolomic data. Left, pathways supported exclusively by transcriptomic data. Right, pathways supported exclusively by metabolomic data.

**Fig. S11.**
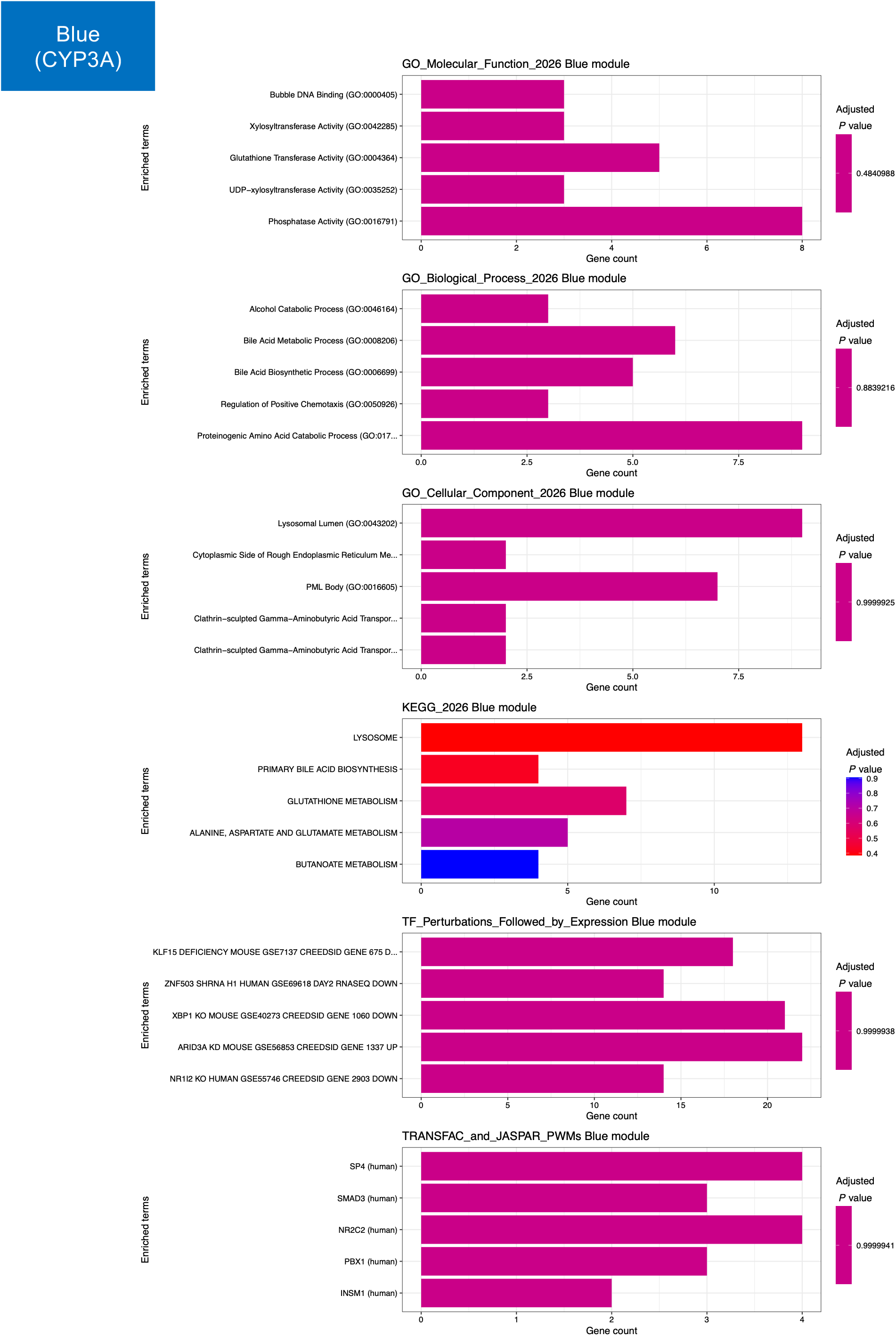
Functional enrichment analysis of genes in the CYP3A module. Rows show Gene Ontology (GO) molecular function, GO biological process, GO cellular component, KEGG pathways, transcription factor perturbation signatures, and TRANSFAC transcription factor motif enrichment analyses.

**Fig. S12.**
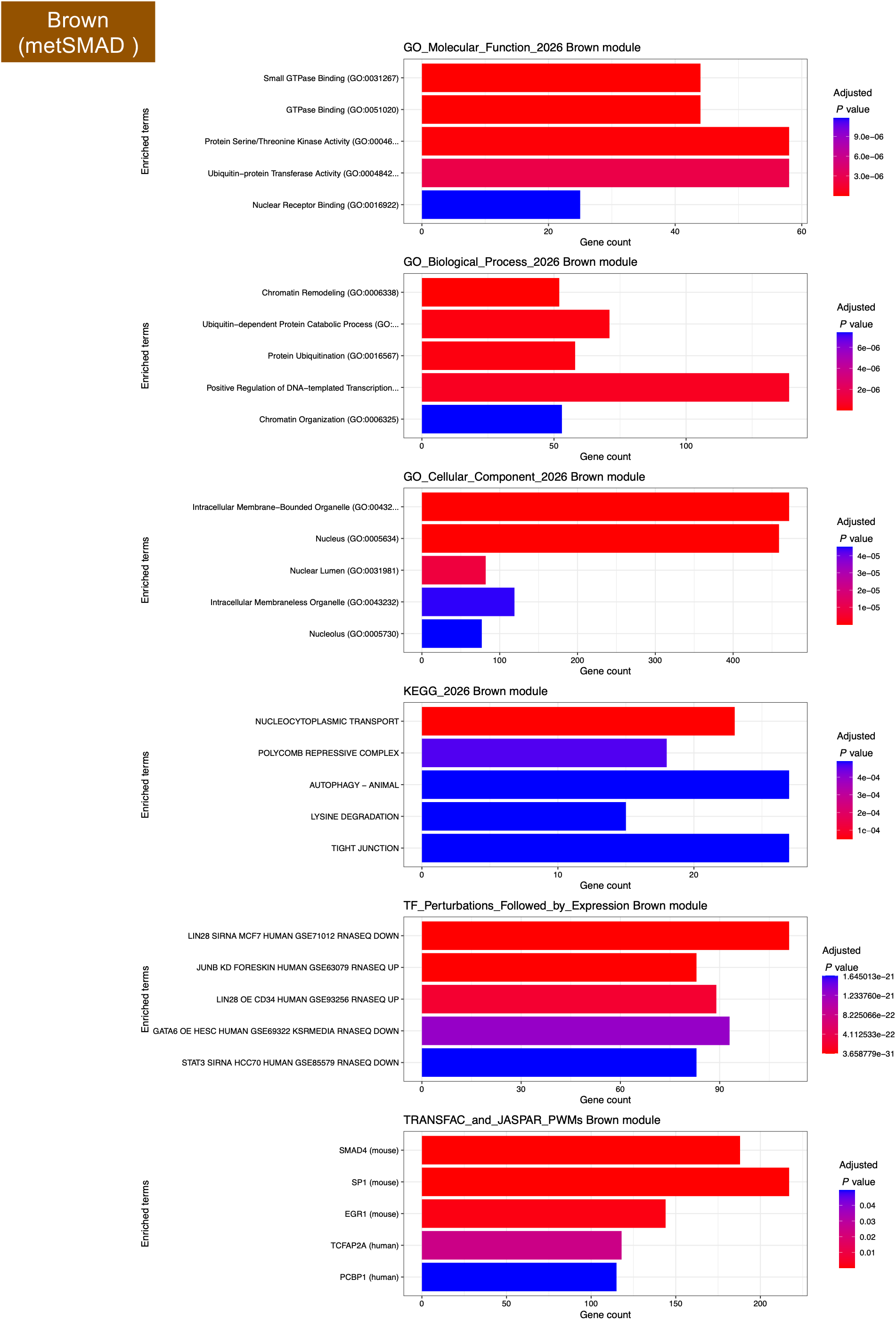
Functional enrichment analysis of genes in the metSMAD module. Rows show Gene Ontology (GO) molecular function, GO biological process, GO cellular component, KEGG pathways, transcription factor perturbation signatures, and TRANSFAC transcription factor motif enrichment analyses.

**Fig. S13.**
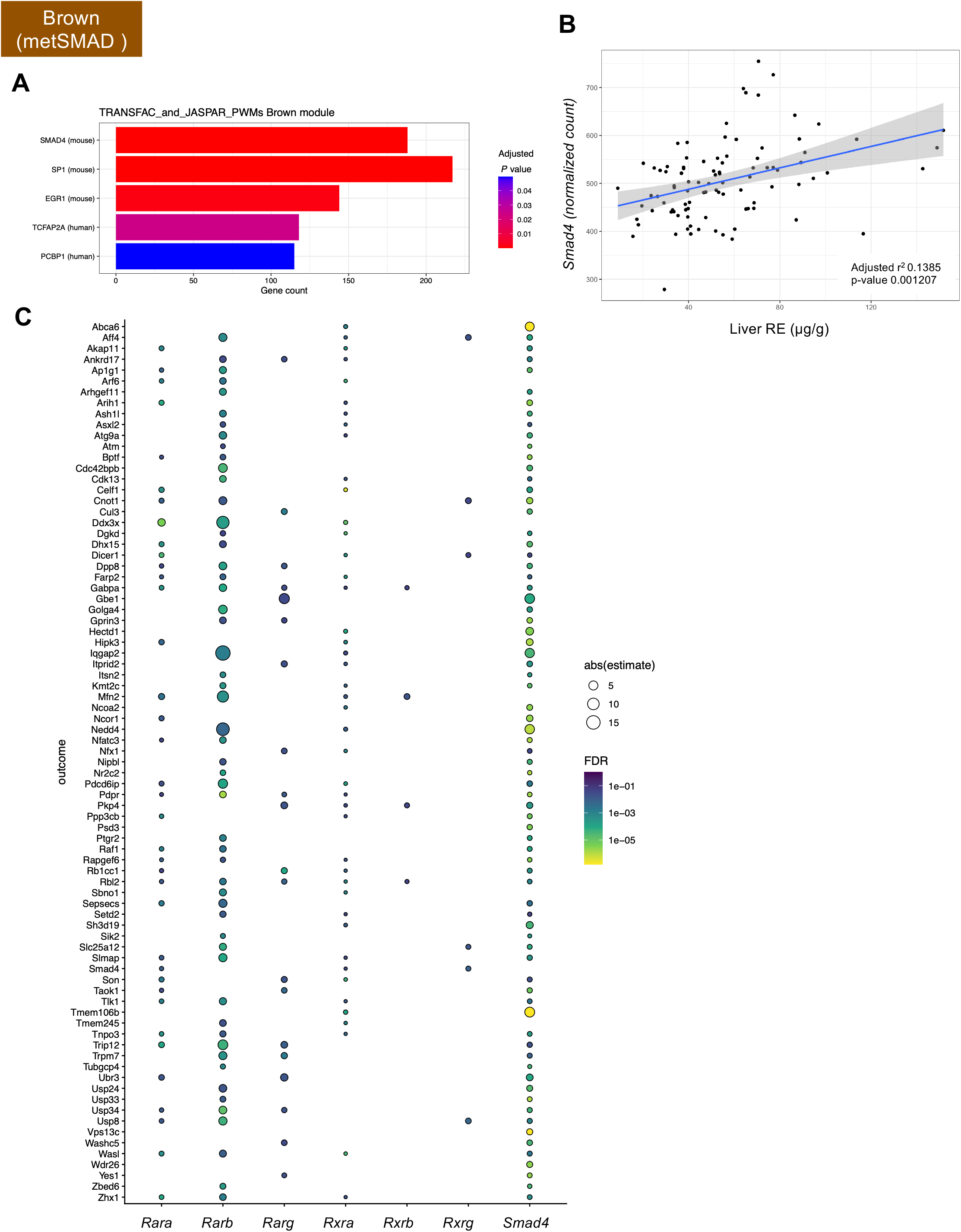
SMAD4 is associated with the metSMAD module and hepatic retinyl ester storage. **(A)** Transcription factor enrichment analysis of metSMAD (brown) module genes. SMAD4-binding motifs were significantly enriched among metSMAD module genes. **(B)** Association between hepatic Smad4 expression and hepatic retinyl ester (RE) concentrations. Higher Smad4 expression was associated with increased hepatic RE levels (adjusted R² = 0.1385, P = 0.001207). **(C)** Association between Smad4 expression and metSMAD module genes. Most highly connected metSMAD module genes were significantly associated with Smad4 expression, consistent with coordinated regulation of the module.

**Fig. S14.**
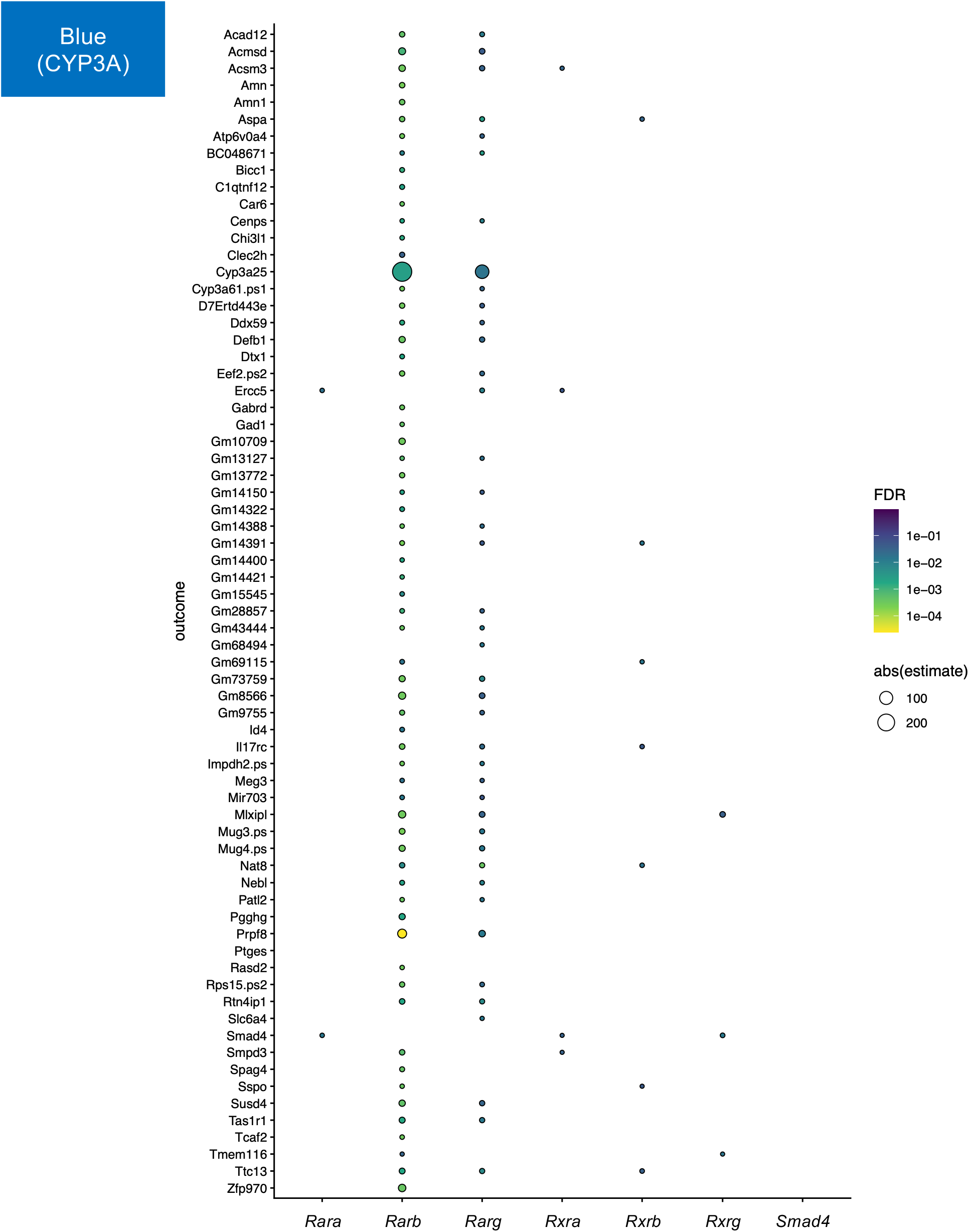
Cyp3a25 is associated with retinoid receptor gene expression within the CYP3A module. Association between expression of the CYP3A module hub gene Cyp3a25 and genes encoding retinoic acid receptors (RARs) and retinoid X receptors (RXRs). Multiple retinoid receptor genes were significantly associated with Cyp3a25 expression.

**Fig. S15.**
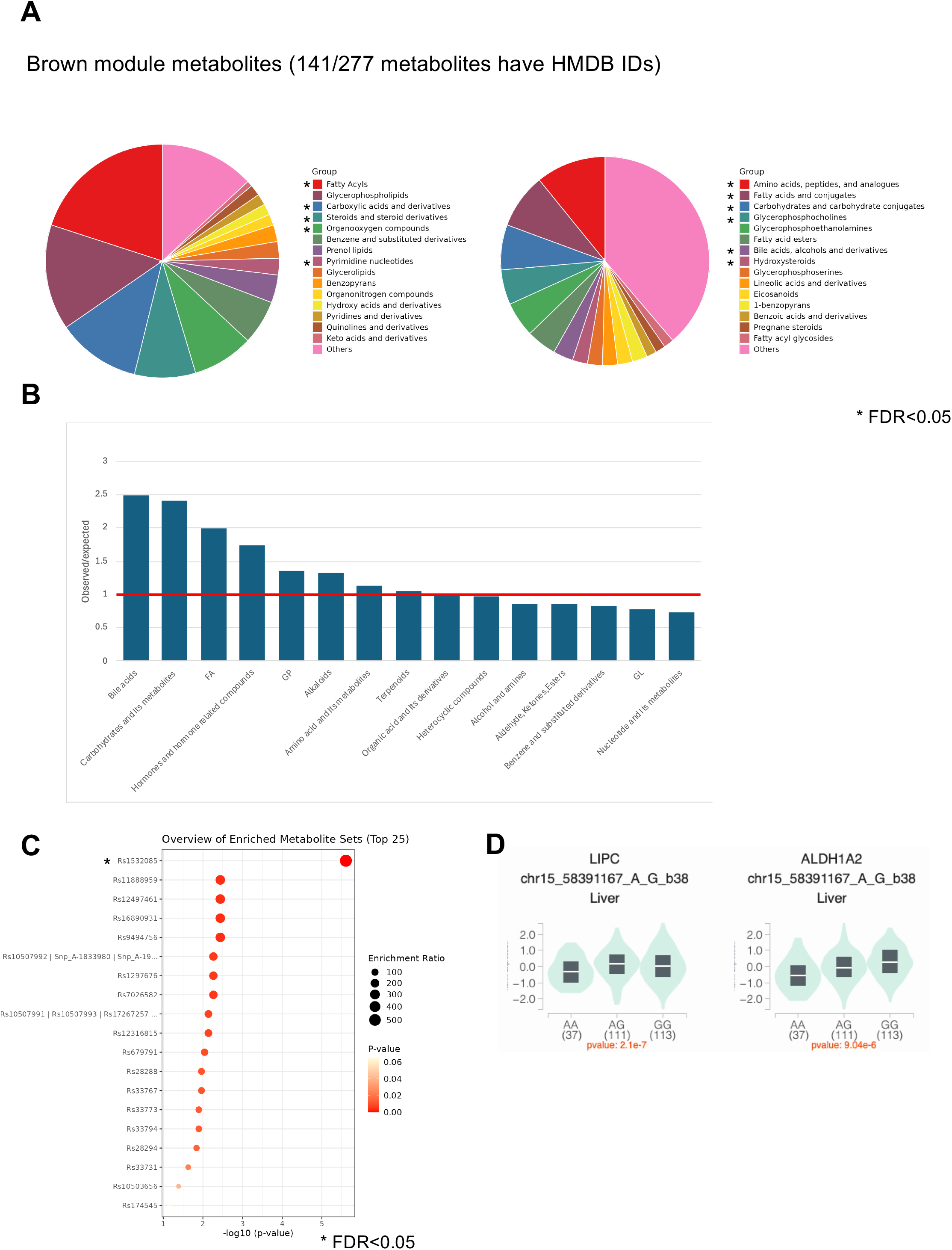
Chemical enrichment analysis of metabolites within the hepatic retinoid-associated brown metabolite module. (A) Enriched metabolite classes identified among metabolites belonging to the brown metabolite module. (B) Enriched metabolite subclasses identified among metabolites belonging to the brown metabolite module. Significant enrichments included amino acids, peptides and analogs; fatty acids and conjugates; carbohydrates and carbohydrate conjugates; and bile acids, alcohols, and derivatives. (C) Enriched Metabolite Sets analysis of metabolites within the brown metabolite module. Metabolites associated with the human single-nucleotide polymorphism (SNP) rs1532085 were significantly enriched. (D) GTEx expression quantitative trait locus (eQTL) data for rs1532085. The rs1532085 genotype was significantly associated with expression levels of *LIPC* and *ALDH1A2*.

**Fig. S16.**
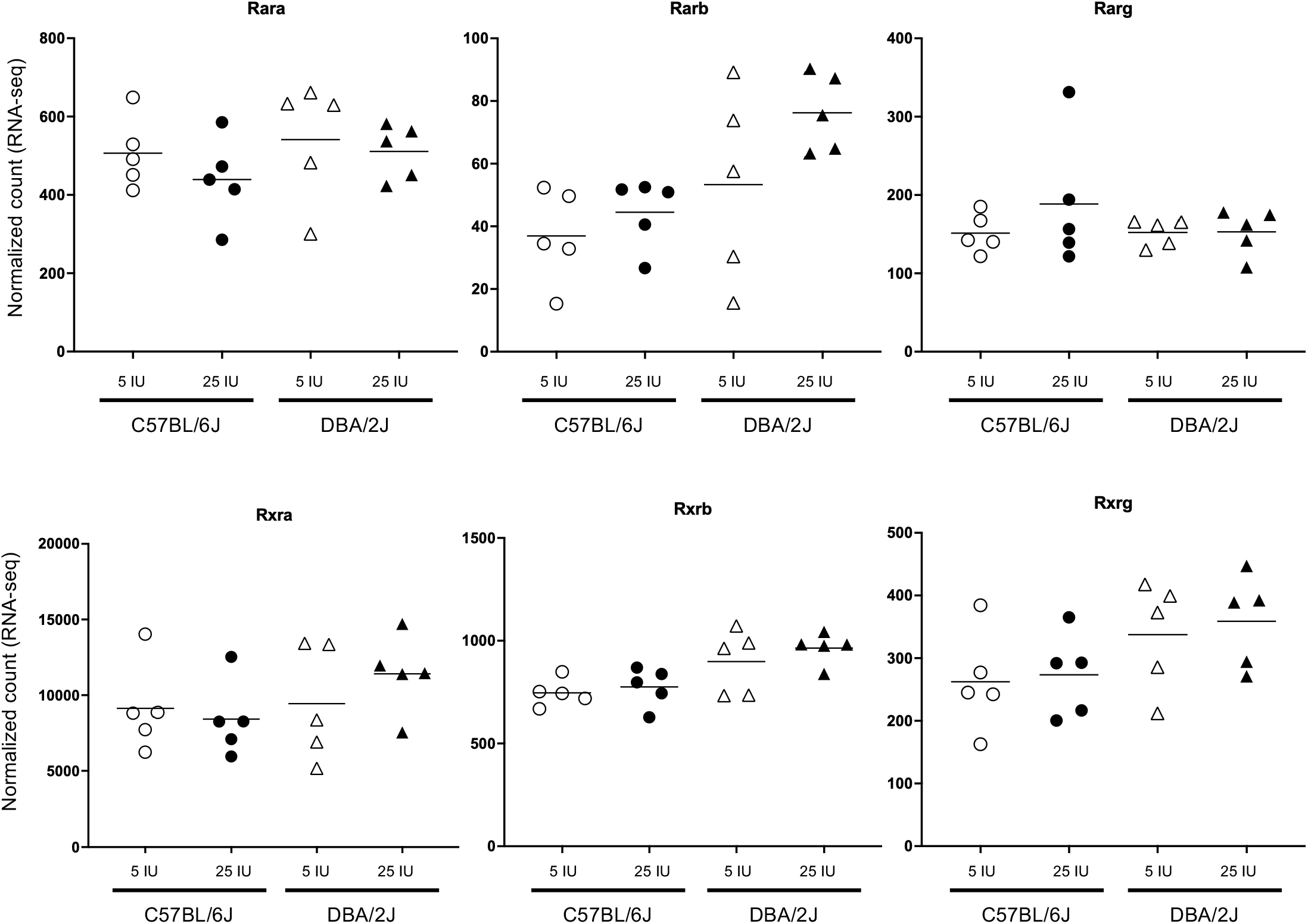
Dietary vitamin A supplementation has minimal effects on hepatic expression of retinoid receptor genes. Normalized RNA-seq counts of *Rara*, *Rarb*, *Rarg*, *Rxra*, *Rxrb*, and *Rxrg* in C57BL/6J and DBA/2J mice fed diets containing 5 or 25 IU vitamin A/g for 10 weeks. Expression of retinoid receptor genes showed little or no response to dietary vitamin A supplementation.

**Fig. S17.**
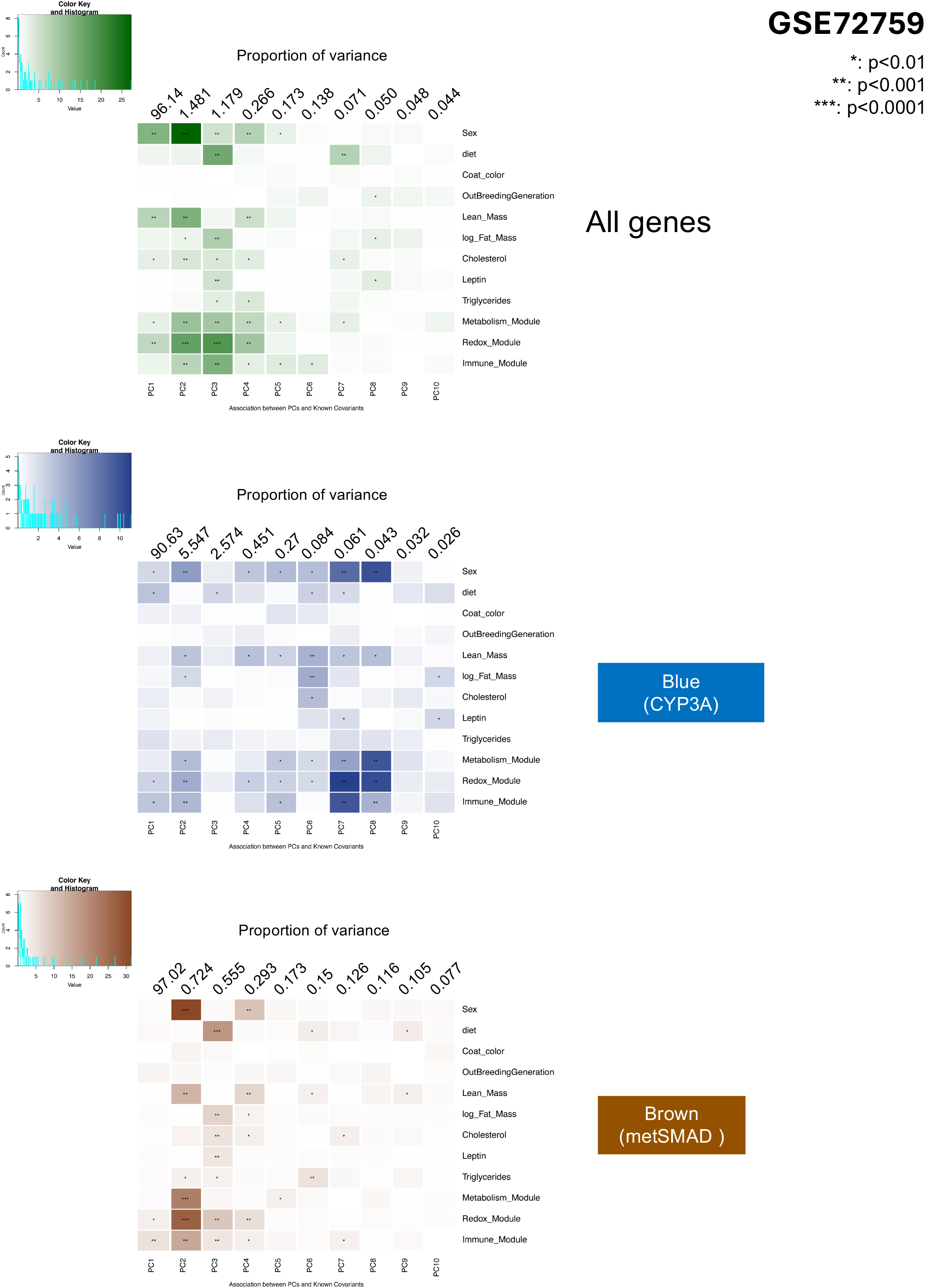
Covariates associated with transcriptional variation in whole-liver and retinoid-associated gene expression profiles in Diversity Outbred mice. **Top**, associations between reported covariates and the first 10 principal components (PCs) derived from genome-wide liver gene expression data (GSE72759). **Middle**, associations between reported covariates and the first 10 PCs derived from CYP3A module hub gene expression. **Bottom**, associations between reported covariates and the first 10 PCs derived from metSMAD module hub gene expression. Colors indicate the significance. *: p<0.01, **: p<0.001, ***: p<0.0001.

**Fig. S18.**
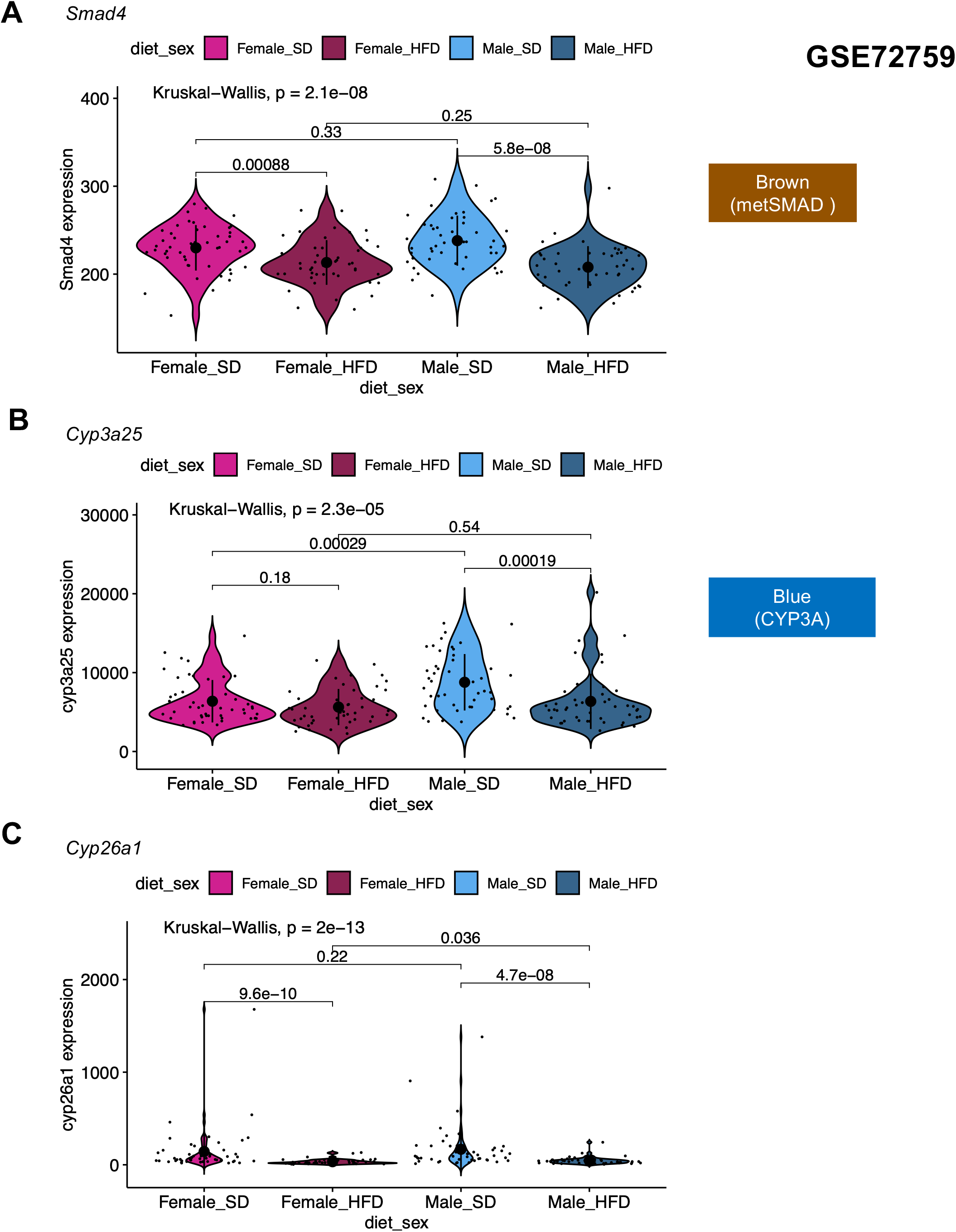
High-fat diet is associated with reduced expression of retinoid-associated hub genes in Diversity Outbred mice. (A-C) RNA-seq counts for Smad4, Cyp3a25, and Cyp26a1 in male and female Diversity Outbred mice maintained on SD or HFD. Expression of all three genes was reduced in HFD-fed animals.

**Fig. S19.**
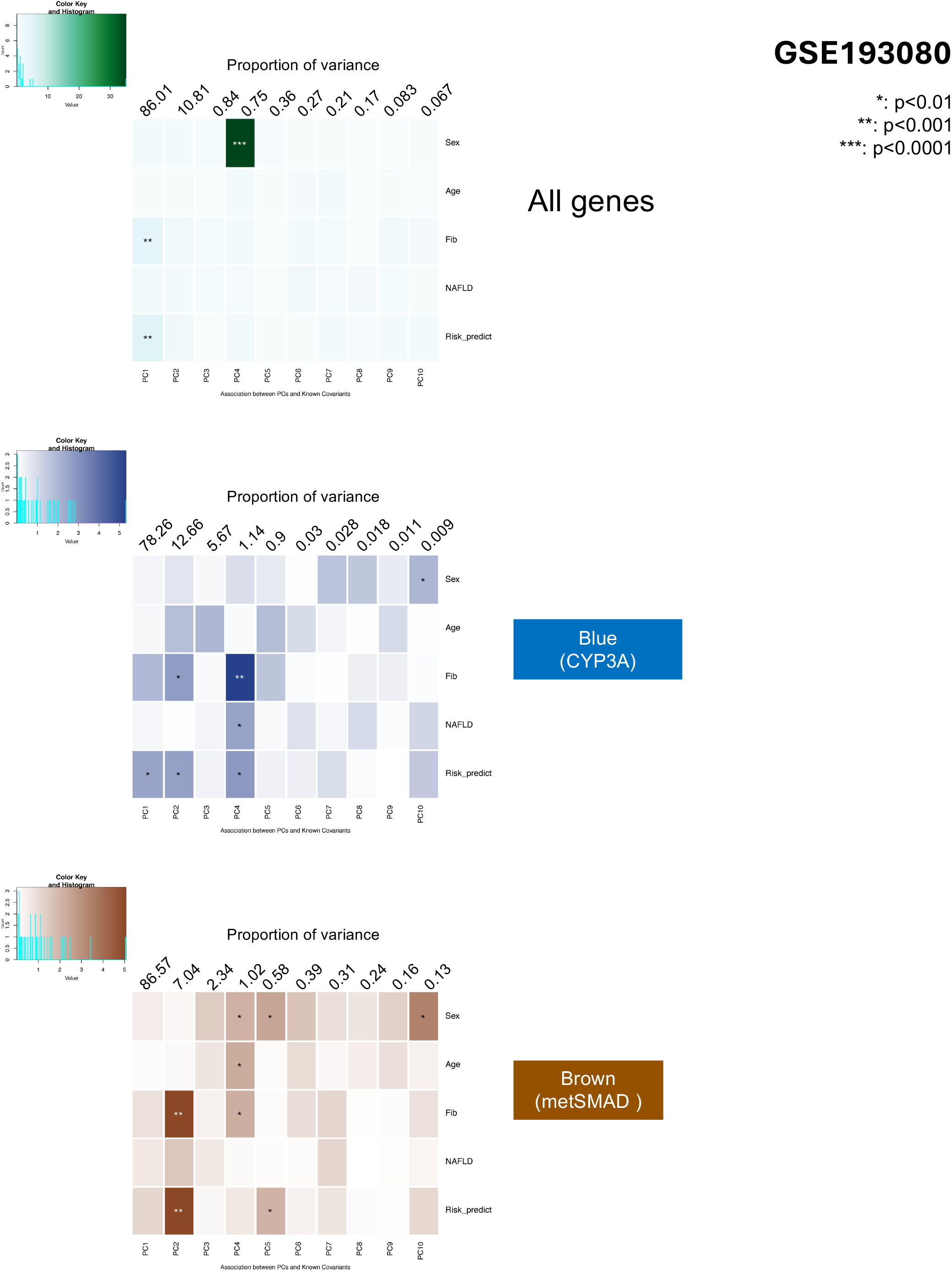
Covariates associated with transcriptional variation in whole-liver and retinoid-associated gene expression profiles in patients with NAFLD (GSE193080). **Top**, associations between reported covariates and the first 10 principal components (PCs) derived from genome-wide liver gene expression data (GSE193080). **Middle**, associations between reported covariates and the first 10 PCs derived from CYP3A module hub gene expression. **Bottom**, associations between reported covariates and the first 10 PCs derived from metSMAD module hub gene expression. Colors indicate the significance. *: p<0.01, **: p<0.001, ***: p<0.0001.

**Fig. S20.**
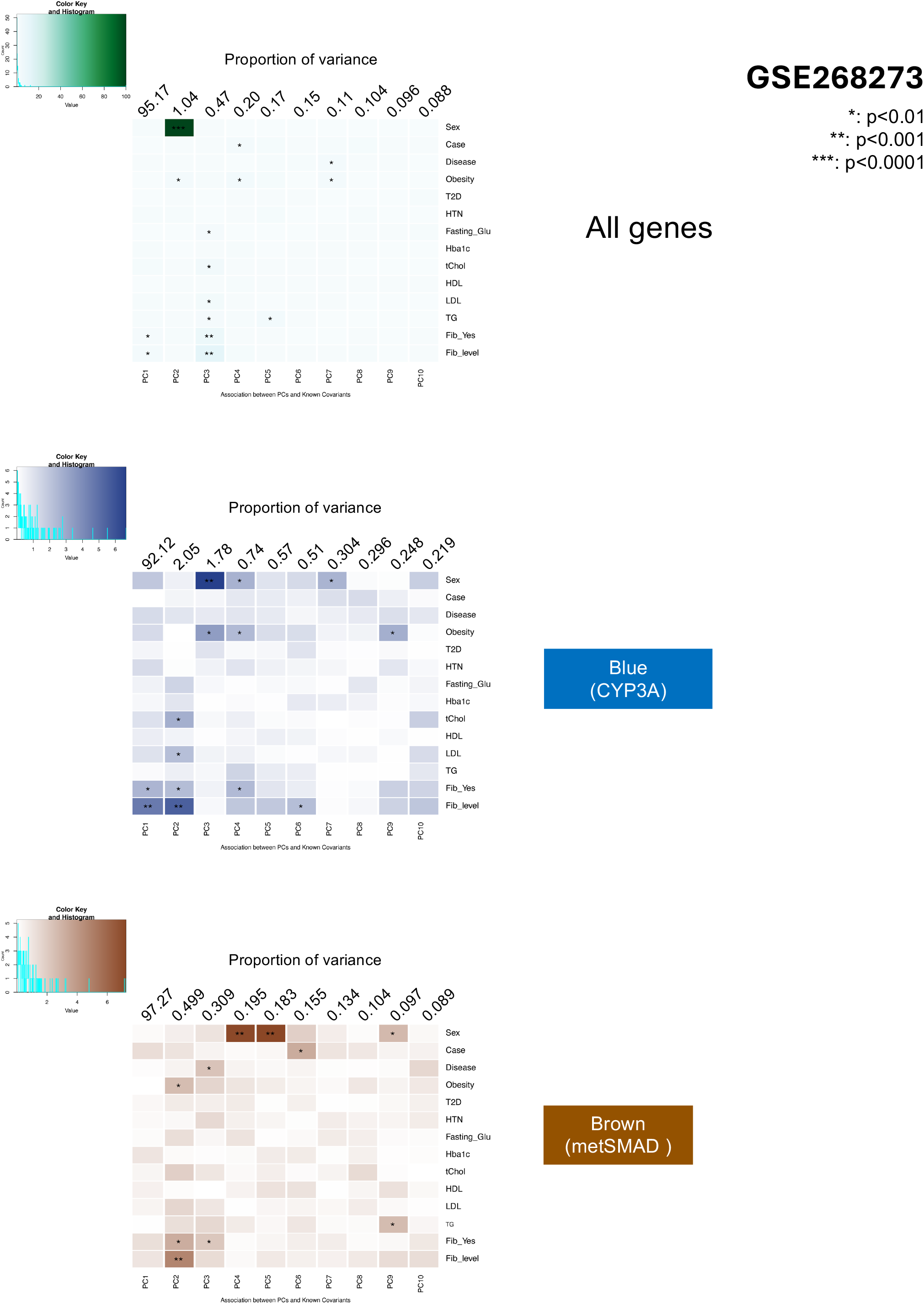
Covariates associated with transcriptional variation in whole-liver and retinoid-associated gene expression profiles in patients with NAFLD and fibrosis (GSE268273). **Top**, associations between reported covariates and the first 10 principal components (PCs) derived from genome-wide liver gene expression data (GSE268273). **Middle**, associations between reported covariates and the first 10 PCs derived from CYP3A module hub gene expression. **Bottom**, associations between reported covariates and the first 10 PCs derived from metSMAD module hub gene expression. Colors indicate the significance. *: p<0.01, **: p<0.001, ***: p<0.0001.

**Fig. S21.**
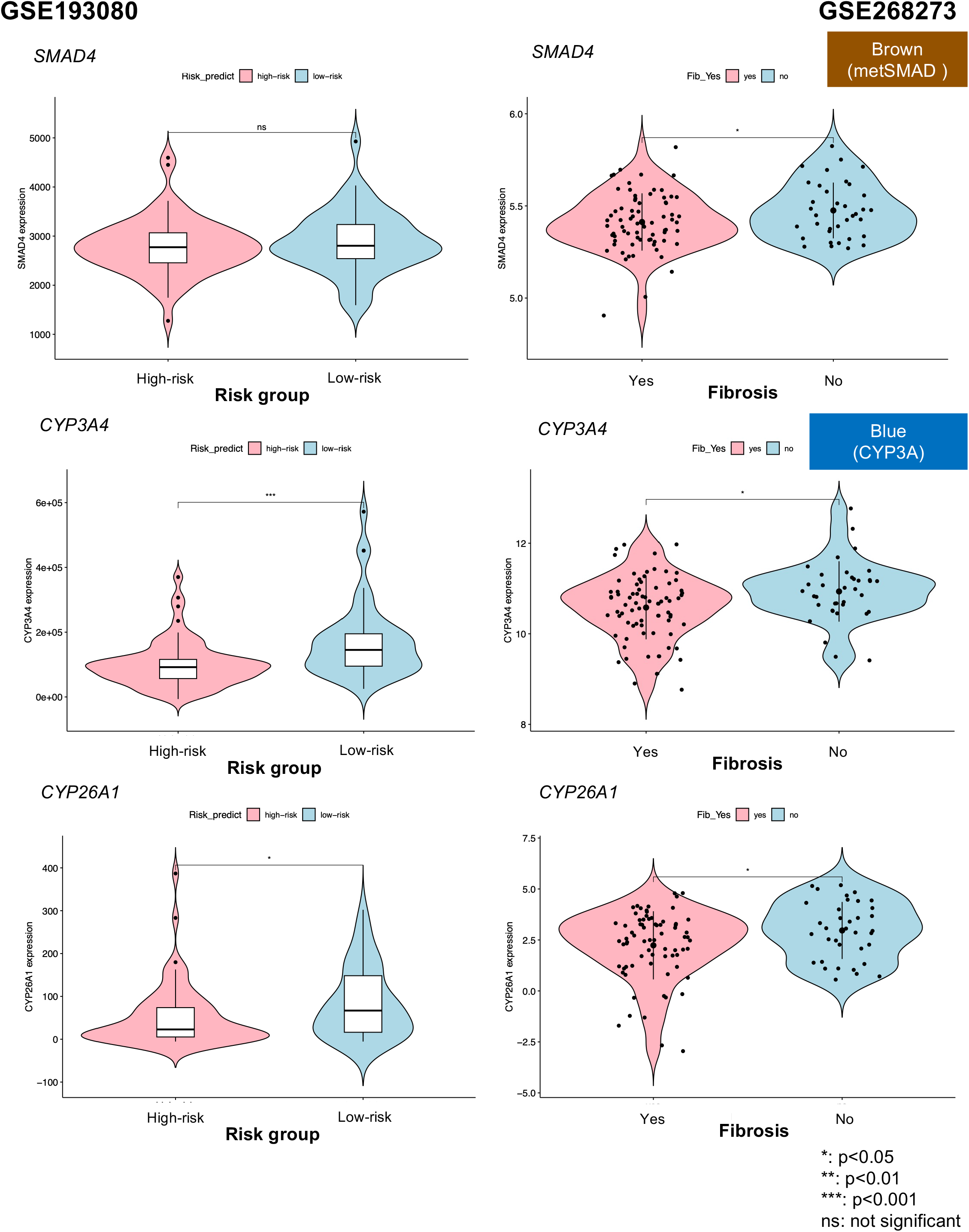
Expression of metSMAD and CYP3A module hub genes in human liver disease transcriptomic datasets. **Left**, GSE193080: comparison of low-risk and high-risk NAFLD patients. **Right**, GSE268273: comparison of patients without fibrosis and with fibrosis. **(A)** Expression of *SMAD4*, the metSMAD module hub gene. **(B)** Expression of *CYP3A4*, the human ortholog of the CYP3A module hub gene *Cyp3a25*. **(C)** Expression of *CYP26A1*, a marker of hepatic retinoid status. *: p<0.05, **: p<0.01, ***: p<0.001.

**Fig. S22.**
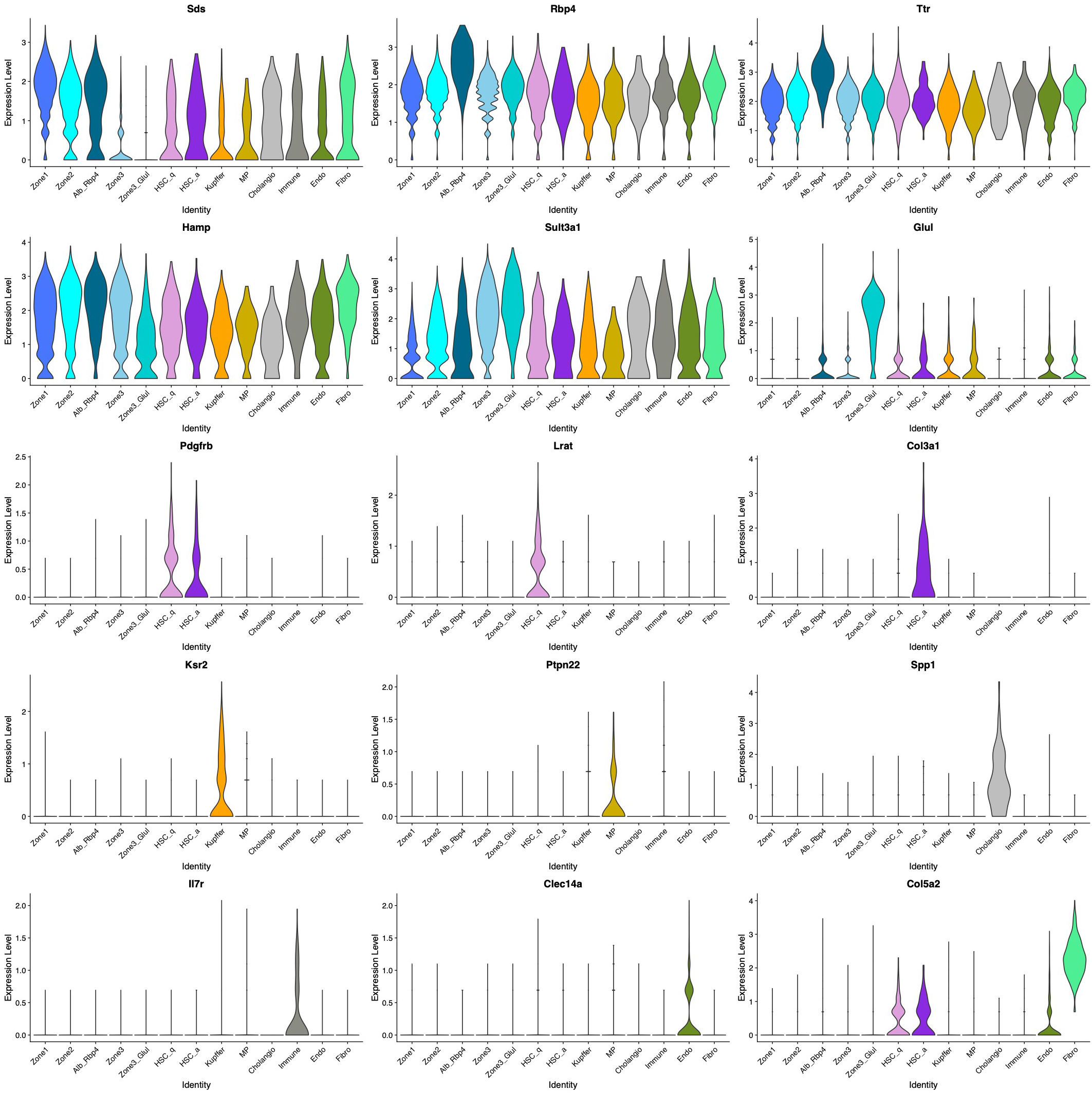
Annotation of liver cell populations by marker gene expression. Violin plots of representative marker genes used for cell-type identification in the snRNA-seq dataset. Expression patterns support the annotation of hepatocyte subpopulations, hepatic stellate cells, Kupffer cells, mononuclear phagocytes, cholangiocytes, immune cells, endothelial cells, and fibroblasts.

**Fig. S23.**
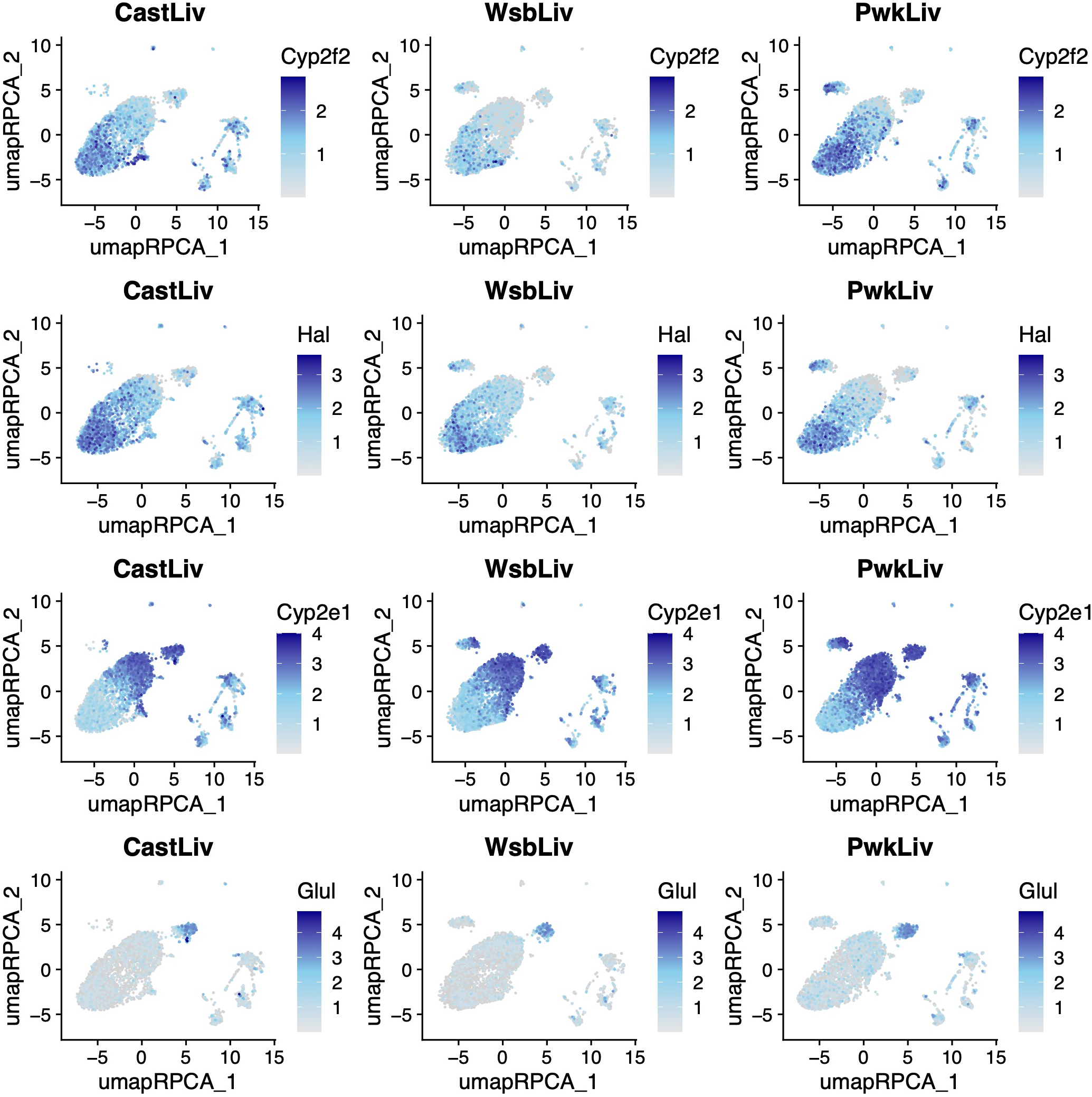
Expression of hepatocyte zonation markers across wild-derived mouse strains. Feature plots showing the expression of representative hepatocyte zonation markers in CAST/EiJ (left), WSB/EiJ (middle), and PWK/PhJ (right) livers. Top row: Cyp2f2 (Zone 1 marker); second row: Hal (Zone 1 marker); third row: Cyp2e1 (Zone 3 marker); bottom row: Glul (pericentral Zone 3 marker). Expression patterns support the identification of zonated hepatocyte populations across all three strains.

**Fig. S24.**
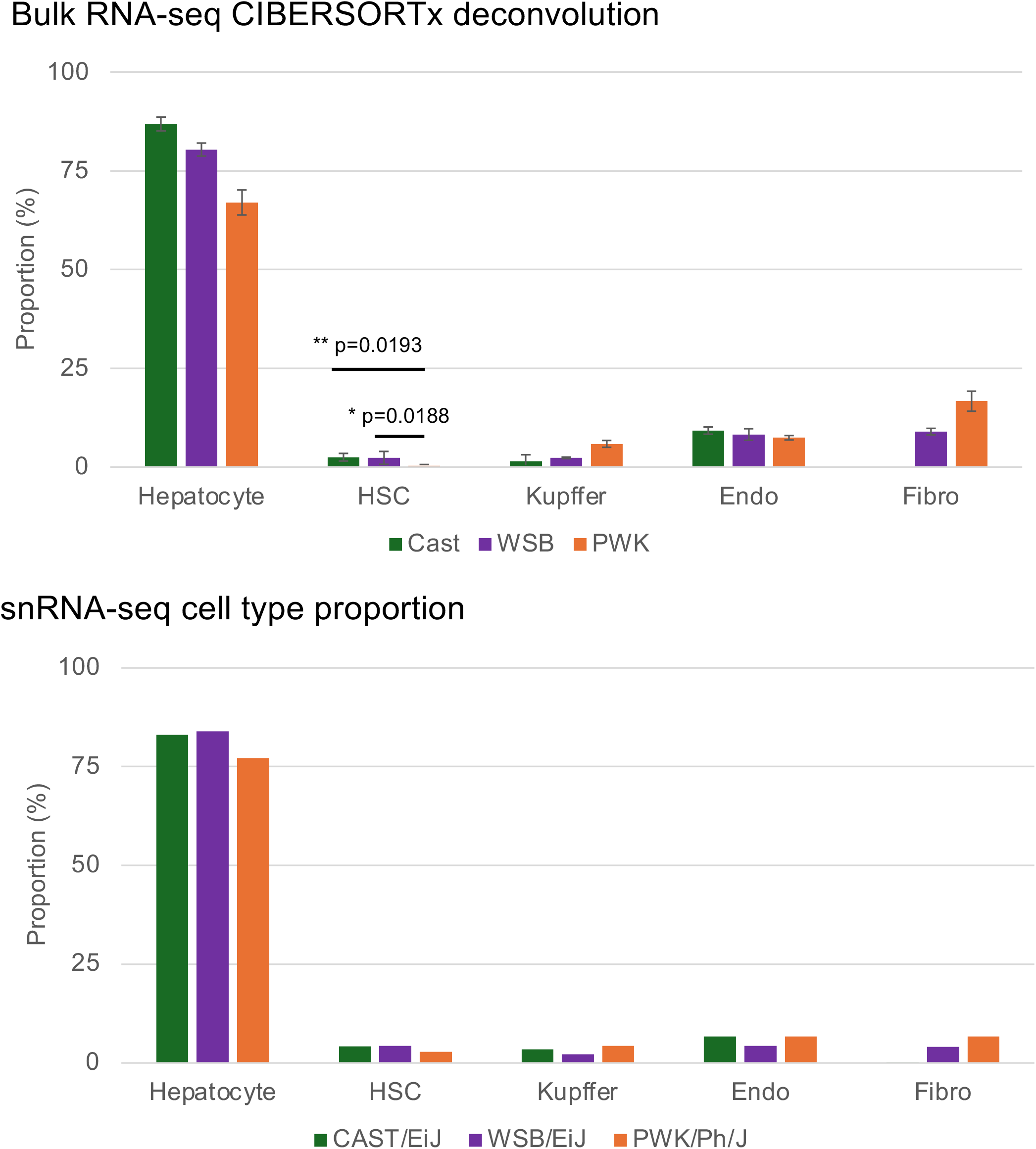
CIBERSORTx deconvolution analysis confirms strain-dependent differences in hepatic stellate cell abundance. **Top**, estimated liver cell-type proportions derived from bulk liver RNA-seq data using CIBERSORTx (n = 5 per strain). **Bottom**, cell-type proportions measured directly by single-nucleus RNA sequencing. Consistent with the snRNA-seq results, PWK/PhJ mice exhibited a lower proportion of hepatic stellate cells than CAST/EiJ and WSB/EiJ mice, supporting strain-dependent differences in liver cellular composition.

**Fig. S25.**
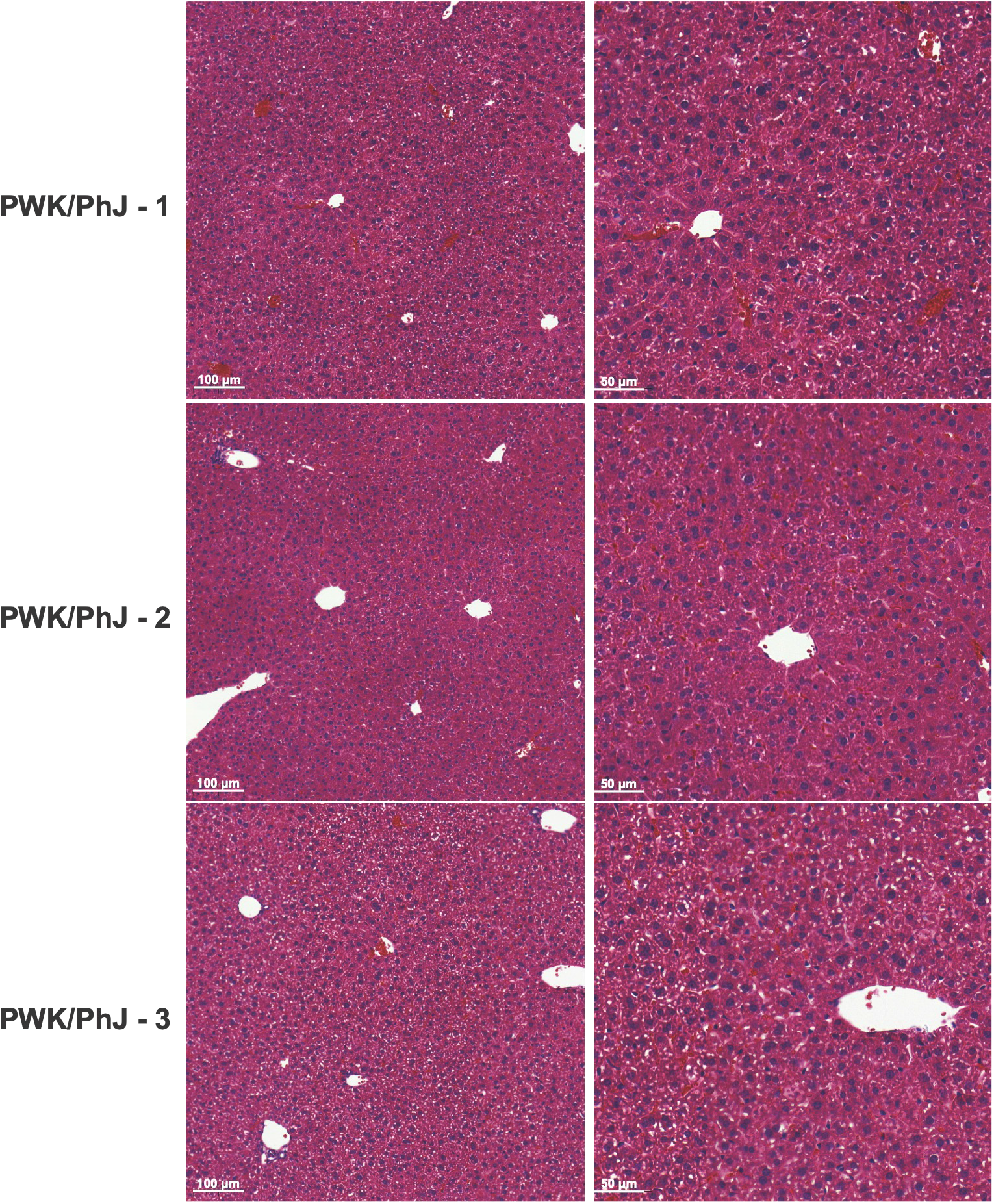
Histopathological assessment of liver tissue from PWK/PhJ mice reveals no overt fibrosis. Representative H&E-stained liver sections from PWK/PhJ mice (n = 5). Histopathological evaluation was performed on all animals, and images from three representative mice are shown. No overt fibrotic lesions were detected. Left, 10× magnification. Right, 20× magnification.

**Fig. S26.**
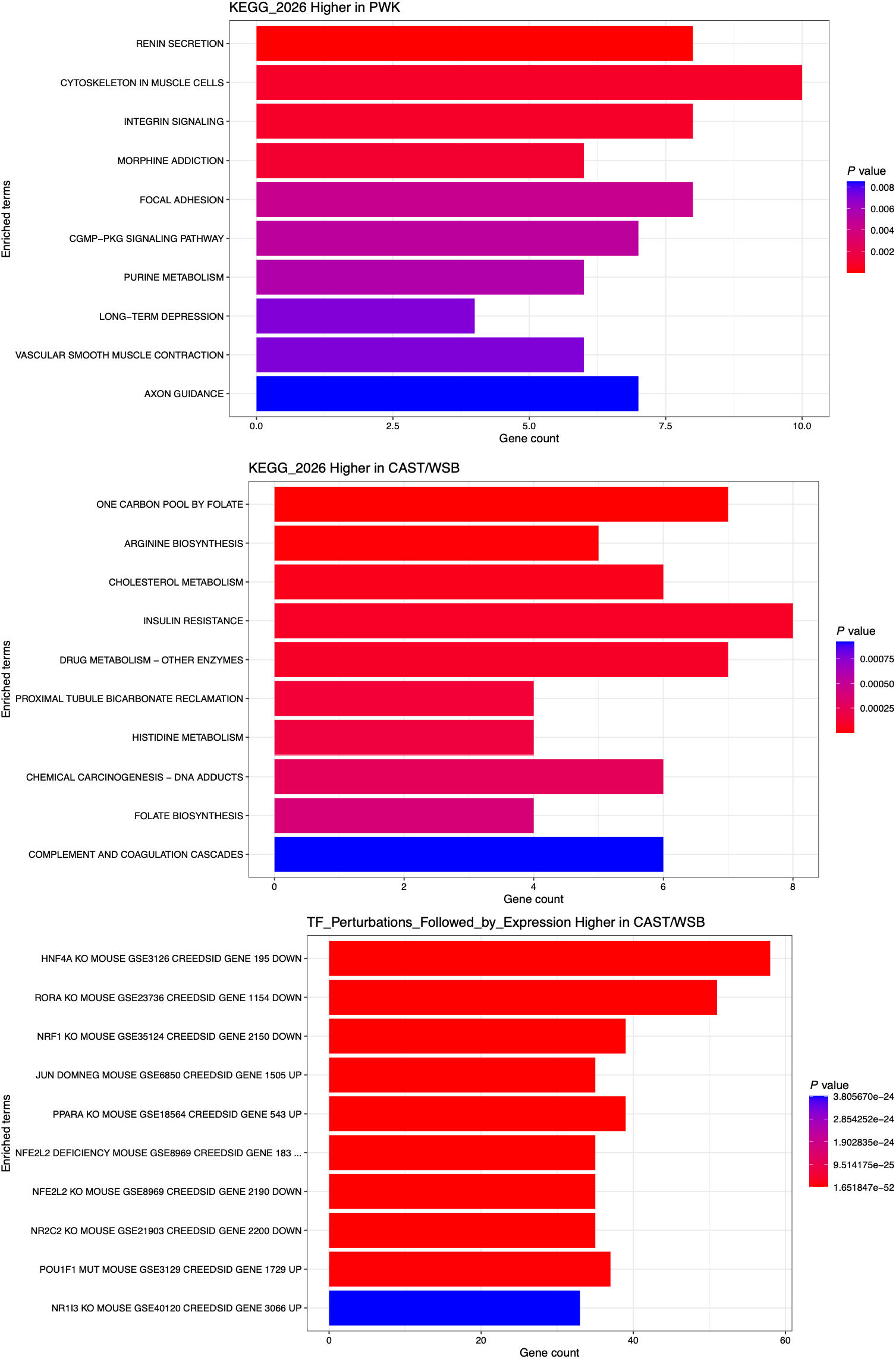
Differential pathway enrichment analysis of hepatic stellate cell transcriptional profiles across wild-derived mouse strains. **Top**: KEGG pathway enrichment analysis of genes expressed at higher levels in PWK/PhJ hepatic stellate cells. Enriched pathways included cytoskeletal organization, integrin signaling, focal adhesion, and vascular smooth muscle contraction. **Middle**: KEGG pathway enrichment analysis of genes expressed at higher levels in CAST/EiJ and WSB/EiJ hepatic stellate cells. Enriched pathways were predominantly associated with metabolic functions. **Bottom**: transcription factor perturbation enrichment analysis of genes expressed at higher levels in CAST/EiJ and WSB/EiJ hepatic stellate cells.

